# ATP-binding cassette protein ABCF1 couples gene transcription with maintenance of genome integrity in embryonic stem cells

**DOI:** 10.1101/2020.05.28.122184

**Authors:** Eun-Bee Choi, Munender Vodnala, Madeleine Zerbato, Jianing Wang, Jaclyn J. Ho, Carla Inouye, Yick W. Fong

**Affiliations:** Brigham Regenerative Medicine Center, Brigham and Women’s Hospital, Boston, MA, USA; Department of Medicine, Cardiovascular Medicine Division, Harvard Medical School, Boston, MA, USA; Department of Molecular and Cell Biology, Li Ka Shing Center for Biomedical and Health Sciences, California Institute for Regenerative Medicine Center of Excellence, University of California at Berkeley, Berkeley, CA, USA; Howard Hughes Medical Institute, Berkeley, CA, USA

**Author notes:** These authors contributed equally (Eun-Bee Choi, Munender Vodnala, Madeleine Zerbato).

## Abstract

OCT4 and SOX2 confer pluripotency by recruiting coactivators to activate stem cell-specific gene expression programs. However, the composition of coactivator complexes and their roles in maintaining stem cell fidelity remain unclear. Here we report the identification of ATP-binding cassette subfamily F member 1 (ABCF1) as a critical coactivator for OCT4/SOX2. ABCF1 is required for pluripotency gene expression and stem cell self-renewal. ABCF1 binds co-dependent coactivators XPC and DKC1 via its intrinsically disordered region and stimulates transcription by linking SOX2 to the transcription machinery. Furthermore, in response to pathogen infection and DNA damage, ABCF1 binds intracellular DNAs accumulated in cells, concomitant with loss of SOX2 interaction and pluripotency gene transcription. This results in spontaneous differentiation of compromised stem cells and elimination from the self-renewing population. Thus, ABCF1 directly couples pluripotency gene transcription with sensing aberrant DNAs and acts as a checkpoint for self-renewal to safeguard stem cell fidelity and genome integrity.

## INTRODUCTION

Stem cell pluripotency is largely driven by core transcription factors including OCT4 and SOX2 (Dunn et al., 2014; Young, 2011). This is exemplified by their ability to induce pluripotency in somatic cells, by reactivating a gene expression program found in embryonic stem (ES) cells (Takahashi and Yamanaka, 2006; Yu et al., 2007). Genome-wide studies demonstrated extensive co-binding of OCT4 and SOX2 at key pluripotency genes and across the ES cell genome (Boyer et al., 2005; Hainer et al., 2019; Kim et al., 2008; Marson et al., 2008). Transcriptional activation of these pluripotency genes by OCT4 and SOX2 requires stem cell-specific coactivators (Fong et al., 2012; Näär et al., 2001; Roeder, 2005). Despite a plethora of these factors already implicated to participate, somatic cell reprogramming remains a highly inefficient and stochastic process, suggesting that additional components may be required for OCT4 and SOX2 to robustly activate pluripotency gene expression (Brumbaugh et al., 2019; Hanna et al., 2009; Jaenisch and Young, 2008; Plath and Lowry, 2011; Takahashi and Yamanaka, 2016). Therefore, there is a need to identify the precise composition of transcriptional complexes assembled at pluripotency gene promoters and the mechanism by which these complexes contribute to stem cell self-renewal.

We took an unbiased in vitro approach to identify coactivators that can reconstitute transcriptional activation by OCT4 and SOX2 (Fong et al., 2011). By biochemical fractionation of human pluripotent cell nuclear extracts and using the well-characterized human *NANOG* promoter as a model transcription template (Kuroda et al., 2005; Rodda et al., 2005), we uncovered three coactivators that are enriched in ES cells and specifically required by OCT4 and SOX2 to activate transcription in vitro. We reported previously that the first two stem cell coactivators (SCCs) are the XPC nucleotide excision repair complex and the dyskerin (DKC1) ribonucleoprotein complex (RNP) (Cattoglio et al., 2015; Fong et al., 2011, 2013, 2014; Zhang et al., 2015). However, we found that optimal activation of the *NANOG* gene by XPC and DKC1 requires an additional coactivator activity (SCCB) (Figure 1). Therefore, revealing the identity and the mechanisms by which SCC-B coordinates with XPC and DKC1 to generate a transcriptional response that promotes the stem cell fate is fundamental to understanding the molecular basis of pluripotency.

**Figure 1.**
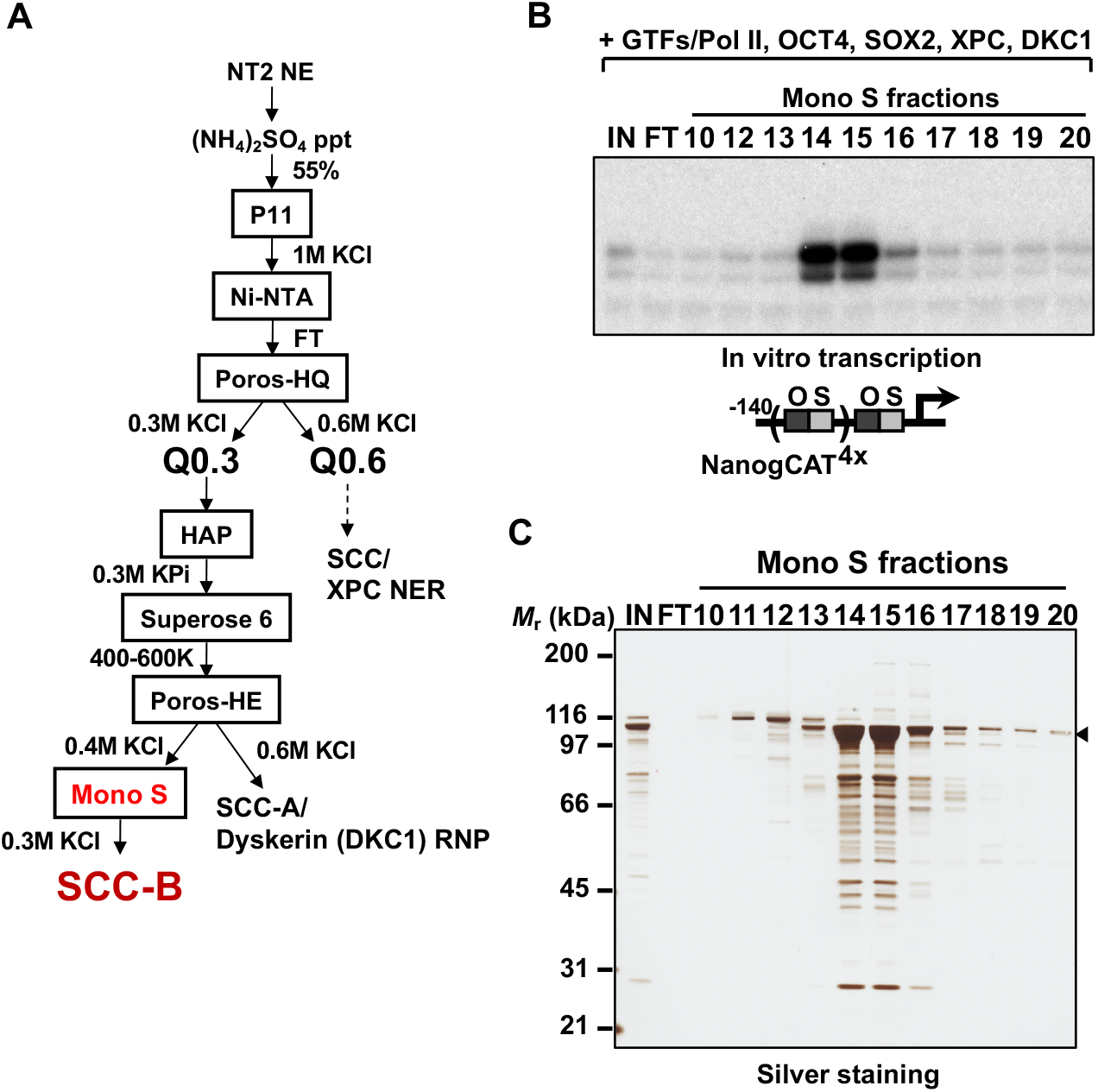
Purification of Stem Cell Coactivator-B (SCC-B). (A) Chromatography scheme for purification of SCC-B from NT2 nuclear extracts (NT2 NE). NT2 NE is first subjected to ammonium sulfate precipitation (55% saturation) followed by a series of chromatographic columns including nickel affinity agarose (Ni-NTA), cation exchangers phosphocellulose (P11), heparin (Poros-HE), Mono S, anion exchanger Poros-HQ, hydroxyapatite (HAP), and gel filtration medium Superose 6. SCC-B activity segregates from SCC-A (Dyskerin (DKC1) RNP) at the Poros-HE step. (B) Input fraction containing SCC-B activity from the Poros-HE step (IN), flow-through (FT) and various salt-eluted Mono S fractions were assayed for their ability to stimulate OCT4/SOX2-dependent transcription from the human *NANOG* promoter template engineered with four extra copies of the oct-sox composite binding element (bottom). All reactions contain purified general transcription factors (GTFs), Pol II, OCT4, SOX2, and recombinant XPC and DKC1 complexes. (C) Fractions assayed in vitro transcription shown in (B) are separated on a 4-12% gradient polyacrylamide gel and stained with silver. Filled arrowhead indicates the predominant polypeptide at ~110 kDa that co-migrates with SCC-B transcriptional activity. The following figure supplement is available for figure 1: Supplementary Figure S1.

Here, we identify SCC-B as ATP-binding cassette subfamily F member 1 (ABCF1). We demonstrate that ABCF1 is required for stem cell maintenance, pluripotency gene expression, and somatic cell reprogramming. Using biochemical approaches and chromatin immunoprecipitation (ChIP) assays, we show that the intrinsically disordered region (IDR) in ABCF1 potently stimulates OCT4/SOX2-dependent transcriptional activation by mediating selective multivalent interactions with XPC, DKC1, SOX2, and RNA polymerase II (Pol II), thereby forming a stem cell-specific transcriptional ensemble at target gene promoters.

In somatic cells, ABCF1 has been implicated in the detection of intracellular DNA and ubiquitin conjugation in the innate immune pathway (Arora et al., 2019; Lee et al., 2013). We provide evidence that leveraging an innate immune DNA sensor such as ABCF1 to regulate pluripotency gene expression may reflect the unique biology of ES cells. Compared to terminally differentiated somatic cells, proliferative ES cells are transcriptionally hyperactive (Efroni et al., 2008). The high replication stress and transcriptional load have been shown to predispose ES cells to DNA damage and genome instability (Fong et al., 2013; Tichy, 2011; Turinetto et al., 2012). In most somatic cell types, the innate immune pathway plays a critical role in managing the DNA damage response (Chatzinikolaou et al., 2014; Harding et al., 2017; MacKenzie et al., 2017; Roers et al., 2016). DNA sensors recognize damage-induced accumulation of endogenous DNA fragments and activate a pro-inflammatory response by stimulating the production of cytokines such as interferons. These cytokines in turn promote the apoptosis and clearance of the affected cells (Jorgensen et al., 2017). Yet, this canonical innate immune pathway is severely attenuated in ES cells and essentially rendered inactive (Guo, 2019). A suppressed innate immune system also suggests that ES cells may lack an effective defense system against pathogen infections. While it is generally thought that pluripotent stem cells are less susceptible to infection because they are physically protected within a blastocyst, vertical transmission of pathogens to these cells can still occur, for example, through infection of the surrounding trophoblast cells, often with devastating consequences to the embryo (Coyne and Lazear, 2016). However, it is clear that these cellular insults must be efficiently resolved, as propagation of rapidly dividing stem cells with compromised genome integrity and cellular fidelity will be deleterious to a developing embryo (Heyer et al., 2000).

We show that ABCF1, a sensor for aberrant intracellular DNA, directly functions as a critical transcriptional coactivator for OCT4 and SOX2. This raises the intriguing possibility of a defense mechanism whereby maintenance of genome integrity in ES cells can be directly linked to pluripotency gene expression by ABCF1. When ES cells are challenged with DNA damage or pathogen-derived DNAs, we show that ABCF1 binds these aberrant DNAs, resulting in loss of interaction with SOX2 and dissociation of ABCF1 from gene promoters targeted by SOX2 and OCT4. This leads to disruption of pluripotency gene expression and elimination of compromised ES cells through spontaneous differentiation (Qin et al., 2007). Our findings reveal that ABCF1 directly couples sensing of infections and genome instability to the pluripotency gene network, thus providing a means for ES cells to maintain a pristine pool of progenitors as they self-renew.

## RESULTS

### A Stem Cell Coactivator Essential for SCC-dependent Transcriptional Activation by OCT4 and SOX2

We have previously shown that robust activation of the human *NANOG* promoter by OCT4 and SOX2 in vitro requires three distinct stem cell coactivators (SCCs) present in a human pluripotent cell nuclear extract (Fong et al., 2011, 2014). The first SCC, the XPC complex, separated from the other two at the Poros-HQ anion exchange chromatographic step, while the second SCC, the DKC1 complex (SCC-A), segregated from the remaining unknown coactivator (SCC-B) at the Poros-Heparin (Poros-HE) step (Figure 1A). Starting with nuclear extracts prepared from 400 L of a pluripotent human embryonal carcinoma cell line N-TERA2 (NT2), we serially fractionated the nuclear extracts over seven chromatographic columns. We tracked SCC-B activity in various protein fractions by assessing their ability to restore SCC-dependent transcriptional activation by OCT4 and SOX2 (Figure S1). In the final Mono S chromatographic step, salt-eluted fractions were assayed in in vitro transcription reactions. We found that adding fractions 14 and 15 to reactions containing purified XPC and DKC1 complexes potently stimulated transcription of the *NANOG* promoter template (Figure 1B). These results demonstrated that SCC-B can dramatically augment the ability of XPC and DKC1 to activate transcription, thus implicating an important role of SCC-B in mediating the cooperative and optimal coactivation by SCCs. Furthermore, our results suggested that the bulk of SCC-B likely resided in these fractions. Accordingly, SDS-PAGE of these Mono S fractions revealed that fractions 14 and 15 were highly enriched with a polypeptide at ~110 kDa along with multiple apparent breakdown products (Figure 1C).

To identify these polypeptides that co-migrate with SCC-B activity, peak Mono S fractions were pooled and separated by SDS-PAGE. Tryptic digestion of the gel slice containing the 110 kDa protein band followed by mass spectrometry identified SCC-B to be ATP-binding cassette subfamily F member 1 (ABCF1) (Figure 2A). We also performed mass spectrometry analysis on the minor protein bands by cutting the gel lane into gel slices and confirmed that they were indeed derived from ABCF1, likely from proteolysis of ABCF1 during biochemical purification (data not shown).

**Figure 2.**
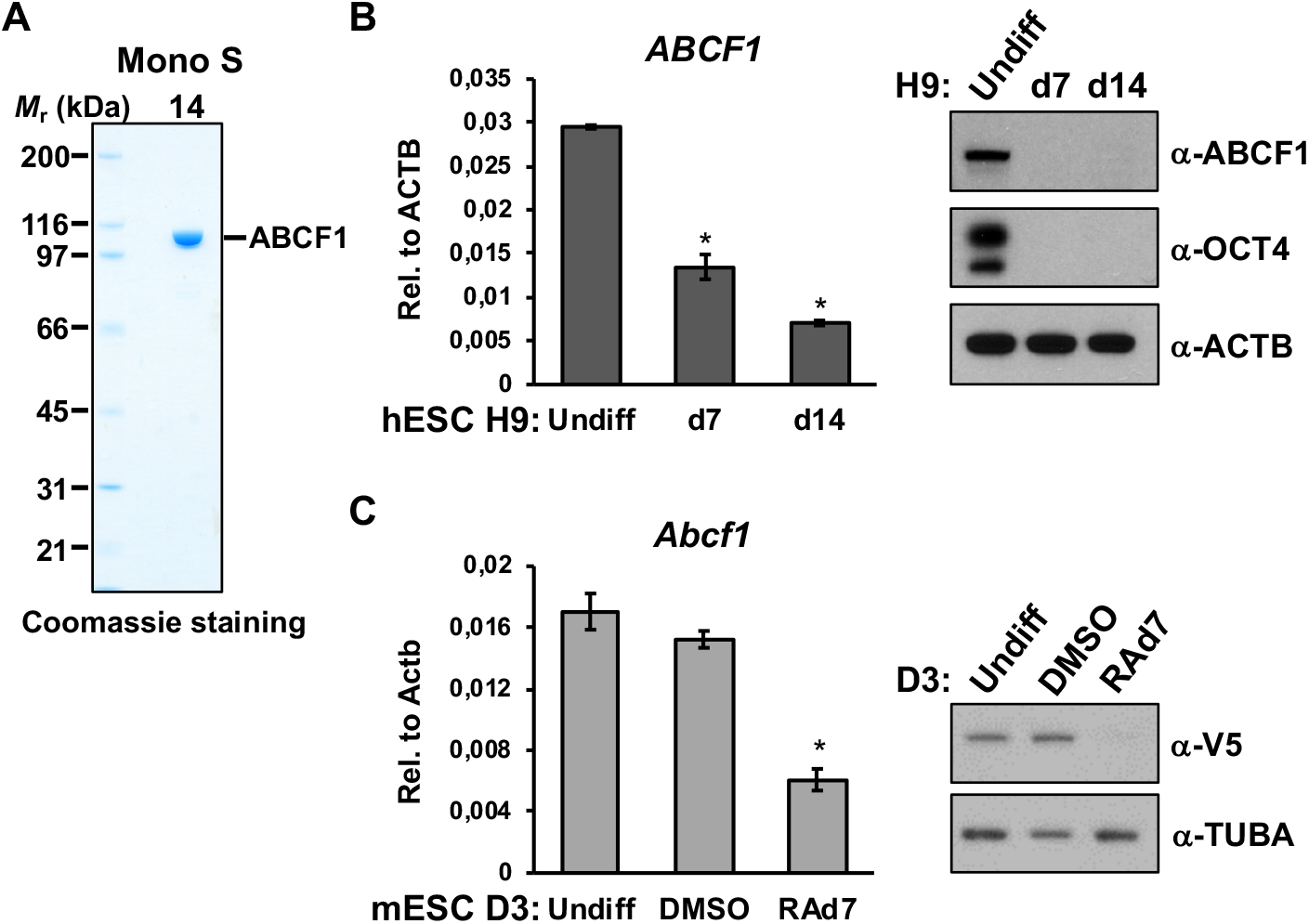
SCC-B is ATP-binding cassette subfamily F member 1 (ABCF1). (A) Coomassie staining of Mono S fraction 14 demonstrates purification to homogeneity. The breakdown products observed in silver staining in Figure 1C are minor compared to the full-length product. Mass spectrometry analysis of Mono S peak activity fractions (#14-16) in Figure 1B identifies the ~110 kDa polypeptide as ABCF1. (B) ABCF1 is enriched in human ES cells. Downregulation of ABCF1 in human ES cell line H9 upon exit of pluripotency. H9 human ES cells are induced to differentiate in differentiation medium (DMEM/F12 with FBS). Cells are collected at day 7 and day 14 post induction. mRNA levels of *ABCF1* are analyzed by quantitative PCR (qPCR) and normalized to *β-actin* (*ACTB*). Western blot analyses of whole cell extracts (WCEs) prepared from H9 cells collected at indicated days are performed using antibodies against ABCF1, pluripotency marker OCT4, and ACTB as loading control. (C) ABCF1 is enriched in mouse ES cells. mRNA and protein levels of ABCF1 are analyzed in pluripotent D3 mouse ES cells carrying V5-epitope tagged *Abcf1* alleles (V5-ABCF1 knock-in), and in the same ES cell line induced to differentiate by retinoic acid (RA) treatment for 7 days (RA d7) by qPCR and western blotting, respectively. Error bars present SEM. *n* = 3. (*) *P* < 0.05, calculated by two-sided Student’s t-test. The following figure supplement is available for figure 2: Supplementary Figure S2.

Identification of ABCF1 as the active constituent of SCC-B activity was unexpected because it has not been previously associated with transcriptional regulation or any cellular processes in the nucleus. To corroborate the mass spectrometry results, we showed by western blotting that ABCF1 resides exclusively in the transcriptionally active phosphocellulose 1M KCl (P1M) nuclear fraction, but not in the inactive P0.3 and P0.5 fractions (Figures 1A and S2) (Fong et al., 2011) Furthermore, we found that the mRNA and protein level of ABCF1 in both human and mouse ES cells decreased sharply when these cells were induced to undergo differentiation (Figures 2B and 2C). Taken together with our previous studies on the expression profile of XPC and DKC1, we showed that all three SCCs are enriched in ES cells but are significantly downregulated upon exit of pluripotency, consistent with these protein factors being deployed to support transcription in a cell type-specific manner (Fong et al., 2011, 2014).

### ABCF1 is Required for Pluripotency Gene Expression and Stem Cell Pluripotency

We next set out to determine if ABCF1 is required for stem cell maintenance and pluripotency by performing loss-of-function studies using lentiviruses expressing shRNAs that target ABCF1 (Figures 3A and S3A). Compared to control D3 mouse ES cells, ABCF1 knockdown resulted in rapid collapse of the tightly packed ES cell colonies and the appearance of flattened and elongated cells with reduced alkaline phosphatase (AP) activity, a marker for pluripotent cells (Figure 3B) (Brambrink et al., 2008; Ginis et al., 2004). We also found that depletion of ABCF1 in both mouse and human ES cells reduced proliferation and viability (Figure S3B). These observations indicated that loss of ABCF1 in ES cells compromises self-renewal capacity and promotes spontaneous differentiation. Consistent with the morphological changes indicative of compromised stem cell identity, depletion of ABCF1 in mouse ES cells resulted in significant decrease in mRNA levels of key pluripotency-associated genes, some of which are known direct targets of OCT4 and SOX2 (Figure 3C) (Chen et al., 2008; Kim et al., 2008; Marson et al., 2008). Concomitant with the downregulation of these pluripotency genes, we observed increased expression of lineage-specific genes related to the ectoderm, mesoderm, and trophectoderm, at the expense of endoderm marker induction (Figure 3D). These data suggest that loss of ABCF1 destabilizes the pluripotency gene network and restricts the ability of ES cells to efficiently differentiate into all three embryonic germ layers. Our finding is consistent with a previously unexplained observation that *Abcf1* knockout mouse embryos die at 3.5 days post coital, a developmental stage that coincides with the emergence of pluripotent cells in the inner cell mass of the blastocyst (Wilcox et al., 2017) and from which ES cells are derived (Brook and Gardner, 1997; Evans and Kaufman, 1981; Martin, 1981).

**Figure 3.**
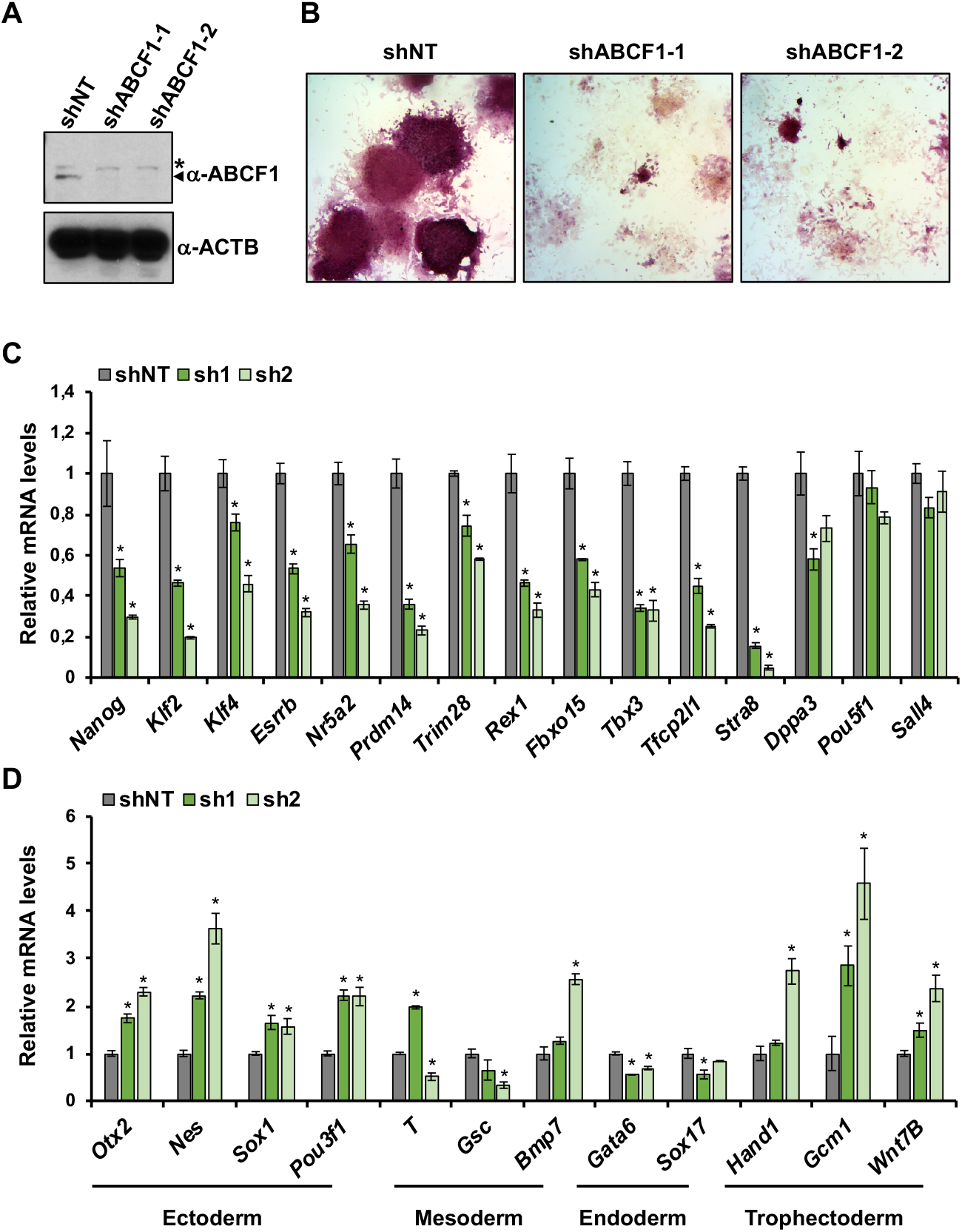
ABCF1 is required for maintenance of stem cell pluripotency. (A) shRNA-mediated knockdown of ABCF1 in mouse ES cells. WCEs of D3 cells transduced with lentiviruses expressing control non-targeting shRNA (shNT) or two independent shRNAs against ABCF1 (shABCF1-1 or shABCF1-2) are analyzed by western blotting to confirm knockdown efficiency. ACTB is used as loading control. Asterisk denotes a non-specific band. (B) shRNA-mediated depletion of ABCF1 in D3 cells leads to colony collapse with flattened cell morphology and reduced alkaline phosphatase (AP) staining, indicating spontaneous differentiation. (C) Loss of ABCF1 in mouse ES cells compromises pluripotency gene expression. Quantification of mRNA levels of pluripotency genes are analyzed by qPCR and normalized to *Actb*. mRNA level in each sample is expressed relative to its respective control set as 1. (D) Depletion of ABCF1 induces expression of differentiation-associated genes. Lineage-specific genes representing the three embryonic germ layers and the trophectoderm are analyzed by qPCR as in (C). Error bars present SEM. *n* = 3. (*) *P* < 0.05, calculated by two-sided Student’s t-test. The following figure supplement is available for figure 3: Supplementary Figure S3.

Given the importance of ABCF1 in stem cell maintenance and expression of genes that are also known to promote somatic cell reprogramming (e.g. *Nanog*, *Klf4*, *Esrrb*, *Prdm14, Nr5a2*) (Figure 3C) (Heng et al., 2010; Iseki et al., 2016; Takahashi and Yamanaka, 2006; Yu et al., 2007), we next assessed whether ABCF1 is required for the generation of induced pluripotent stem (iPS) cells from mouse embryonic fibroblasts (MEFs). To this end, we transduced MEFs with lentiviruses expressing non-targeting control shRNA or two independent shRNAs specific for ABCF1 and initiated reprogramming by doxycycline (dox)-induced expression of OCT4, KLF4, SOX2, and c-MYC (Sommer et al., 2009). Similar to results we previously observed with depletion of XPC and DKC1, ABCF1 knockdown resulted in a marked decrease in the number of AP-positive iPS cell colonies formed (Figure S3C) (Fong et al., 2011, 2014).

Flow cytometry analysis further demonstrated that loss of ABCF1 led to a decrease in cells expressing SSEA-1+, a cell While the significance of cell proliferation rate for somatic cell reprogramming remains unclear (Guo et al., 2014; Gupta et al., 2015; Ruiz et al., 2011; Son et al., 2013; Xu et al., 2013), we should note that ABCF1 knockdown MEFs displayed noticeable changes in growth rate (data not shown). Our results are consistent with a role of ABCF1 in overcoming early barriers to reacquisition of pluripotency during cellular reprogramming.

We reasoned that if indeed ABCF1 functions as a transcriptional coactivator for OCT4 and SOX2 in ES cells, ABCF1 ought to occupy the genome at cis-regulatory regions that are bound by OCT4 and SOX2. Therefore, we surface marker that indicates early stage of reprogramming (Figure S3D) (Brambrink et al., 2008; Polo et al., 2012). performed chromatin immunoprecipitation (ChIP) assays to investigate the binding of ABCF1 at known pluripotency gene loci targeted by OCT4 and SOX2 (Chen et al., 2008; Hainer et al., 2019; Marson et al., 2008). Because we could not find a commercially available antibody against ABCF1 suitable for ChIP, we employed CRISPR/Cas9 genome editing to knock-in a V5 epitope tag into the N-terminus of the *Abcf1* gene to enable ChIP with an anti-V5 antibody. We confirmed D3 mouse ES cell clones with biallelic knock-in of the V5-tag by genomic sequencing and western blotting (Figure S4A). Importantly, these knock-in lines express pluripotency and differentiation-associated genes at a level comparable to wild-type cells, indicating that epitope tagging did not compromise ABCF1 function (Figure S4B). We performed ChIP using formaldehyde as well as the protein-protein crosslinker ethylene glycol bis[succinimidylsuccinate] (EGS) (Zeng et al., 2006), and found that ABCF1 was enriched at OCT4/SOX2 co-bound regions in pluripotency genes such as *Oct4 (Pou5f1)*, *Sox2*, *Lefty1*, and *Trim28* (Figures 4A and 4B). By contrast, we did not observe any significant enrichment of ABCF1 at housekeeping genes *β-actin* (*Actb*) (Figure 4C) and *Dhfr* (Figure S4C), consistent with ABCF1 acting as a selective coactivator for OCT4 and SOX2. Using an alternative ChIP approach that can better preserve antibody epitopes than shearing by sonication, ChIP of crosslinked chromatin digested with micrococcal nuclease (MNase) revealed that ABCF1 was also enriched at the regulatory region of *Nanog* in addition of *Oct4* and *Sox2*, but not *Actb* (Figures 4D and S4D-E). Taken together, the data presented thus far suggest a classical coactivator function of ABCF1 both in vitro with naked DNA and in the context of chromatin in ES cells.

**Figure 4.**
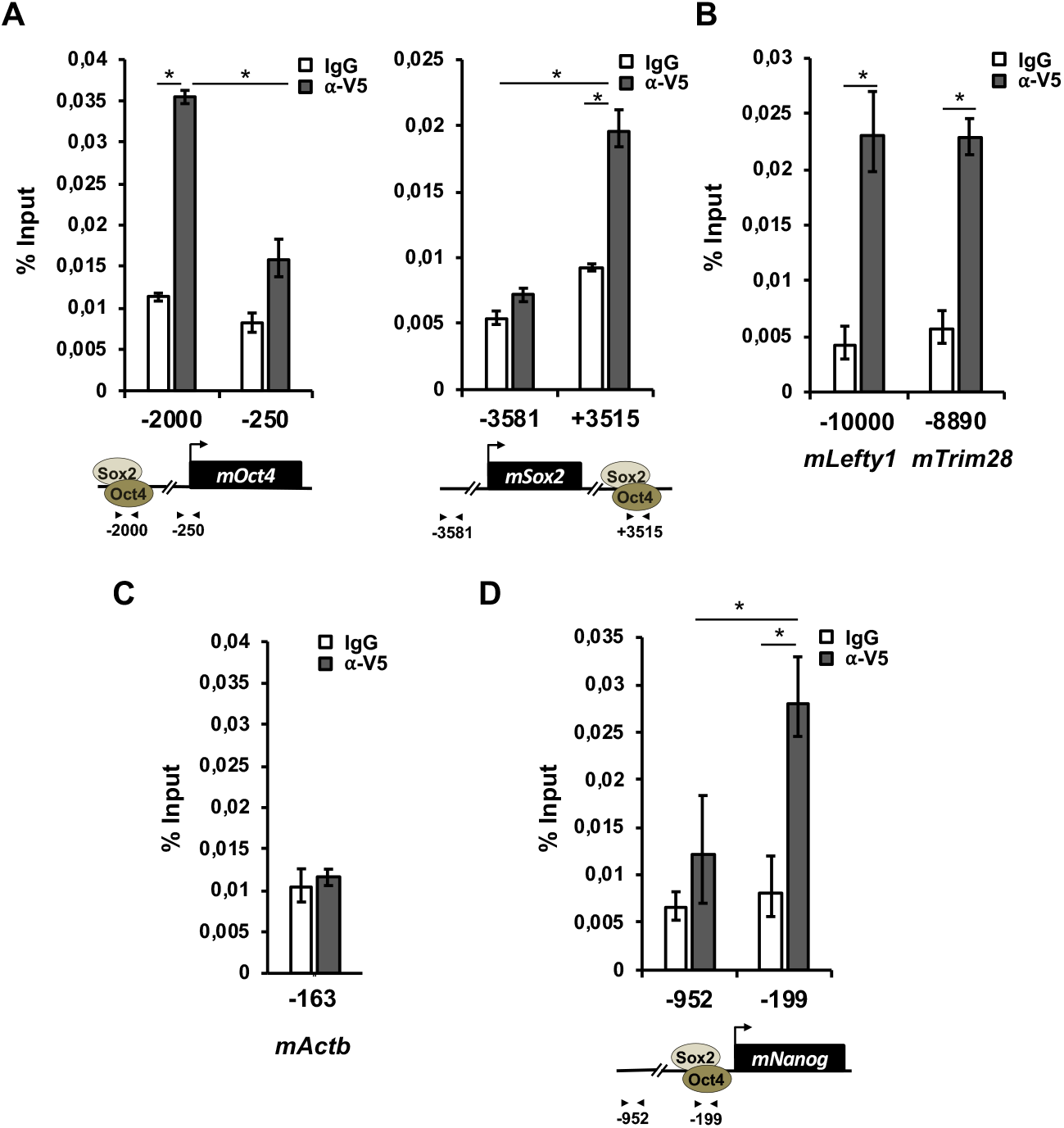
ABCF1 is recruited to regulatory regions of key pluripotency genes targeted by OCT4 and SOX2 in mouse ES cells. (A) Chromatin immunoprecipitation (ChIP) analysis of ABCF1 occupancy on control and enhancer regions of *Oct4* and *Sox2* gene loci in V5-ABCF1 knock-in D3 mouse ES cells. Representative data showing the enrichment of V5-ABCF1 (grey bars) compared to control IgGs (white bars) are analyzed by qPCR and expressed as percentage of input chromatin. (B) ABCF1 is recruited to strong SOX2-bound genomic regions. ABCF1 enrichment at the enhancer regions of *Lefty1* and *Trim28* genes is analyzed as described in (A). (C) ABCF1 is not recruited to the promoter of housekeeping gene *Actb*. Enrichment of ABCF1 is analyzed as in (A). (D) MNase-ChIP analysis of ABCF1 occupancy on the *Nanog* gene promoter. Enrichment of ABCF1 is analyzed as in (A). Schematic diagrams of OCT4/SOX2 binding sites of each gene and the relative positions of the amplicons are shown at the bottom. Error bars present SEM. *n* = 3. (*) *P* < 0.05, calculated by two-sided Student’s t-test. The following figure supplement is available for figure 4: Supplementary Figure S4

### Transcriptional Coactivation Mediated by the Intrinsically Disordered Region in ABCF1

ABCF1 belongs to a large class of transporters that couples ATP hydrolysis to the active transport of substrates across cell membranes (Vasiliou et al., 2009). While most ABC proteins contain transmembrane domains (TMDs) as expected, ABCF1 lacks TMD and is therefore not predicted to function as a transporter. In somatic cells, ABCF1 has been implicated as a regulator of mRNA translation (Coots et al., 2017; Paytubi et al., 2008), an E2 ubiquitin-conjugating enzyme (Arora et al., 2019), and a sensor for intracellular double-stranded DNAs involved in innate immune responses (Lee et al., 2013). It is worth noting that these reported activities of ABCF1 all reside in the cytoplasm. Our findings here thus point to a hitherto unknown nuclear function of ABCF1 in gene regulation in ES cells.

To gain mechanistic insights into how ABCF1 stimulates SCC-dependent transcriptional activation by OCT4 and SOX2, we next sought to delineate the protein domain and catalytic activity (i.e. ATPase) of ABCF1 that are critical for coactivator function. ABCF1 contains two conserved nucleotide-binding domains (NBDs) and a ~ 300 amino acid N-terminal domain that is predicted to be highly disordered (Abor et al., 2018) (Figure S5A). This is consistent with a recent structural study on ABCF1 which indicated that the N-terminal domain cannot be crystallized (Qu et al., 2018). The N-terminal domain of ABCF1 is a low-complexity region that is unusually rich in charged amino acids, of which ~40% are divided between lysine (K) and glutamic acid (E) residues (Dyson, 2016; Hansen et al., 2006; Paytubi et al., 2009) (Figure 5A). This unique amino acid composition is conserved among vertebrates (Figure S5B) and consistent with one of the largest classes of intrinsically disordered region (IDR) called polyampholytes (Das and Pappu, 2013; Van Der Lee et al., 2014). This putative IDR also contains a polyglutamine (polyQ) tract that is found in transactivation domain of many transcription factors (Atanesyan et al., 2012; Zhang and Tjian, 2017). We purified recombinant full-length wild-type (WT) and an ATPase-defective mutant ABCF1 wherein the two conserved ATP-binding lysine residues (K324 and K664) in the Walker A motif of both NBDs were substituted with methionine (2KM) (Paytubi et al., 2009) (Figures 5B and S5C). We also generated the IDR fragment (amino acids 1-302) as well as a series of N-terminal truncations of the IDR in full-length ABCF1. When added to in vitro transcription assay, the full-length WT and 2KM mutant proteins were as active as endogenous ABCF1 purified from NT2 cells (Figure 5C). However, deletion of polyQ alone (Δ115) or together with the K/E-rich domain (Δ248) led to a progressive loss of transcriptional activity, and the two NBDs by themselves (Δ302) were completely inactive. Surprisingly, the IDR fragment (1-302) also lacked transcriptional activity. These results provide several insights. First, ABCF1 is unambiguously shown to be the sole constituent of SCC-B. Second, ATP binding and hydrolysis by ABCF1 are dispensable for transcription, unlike for ABCF1’s role in translation (Paytubi et al., 2009; Qu et al., 2018). Third, while the entire IDR is essential for full coactivator activity, the IDR fragment by itself is not sufficient to activate transcription. These observations suggest that the NBDs also contribute to the full transcriptional activity of ABCF1.

**Figure 5.**
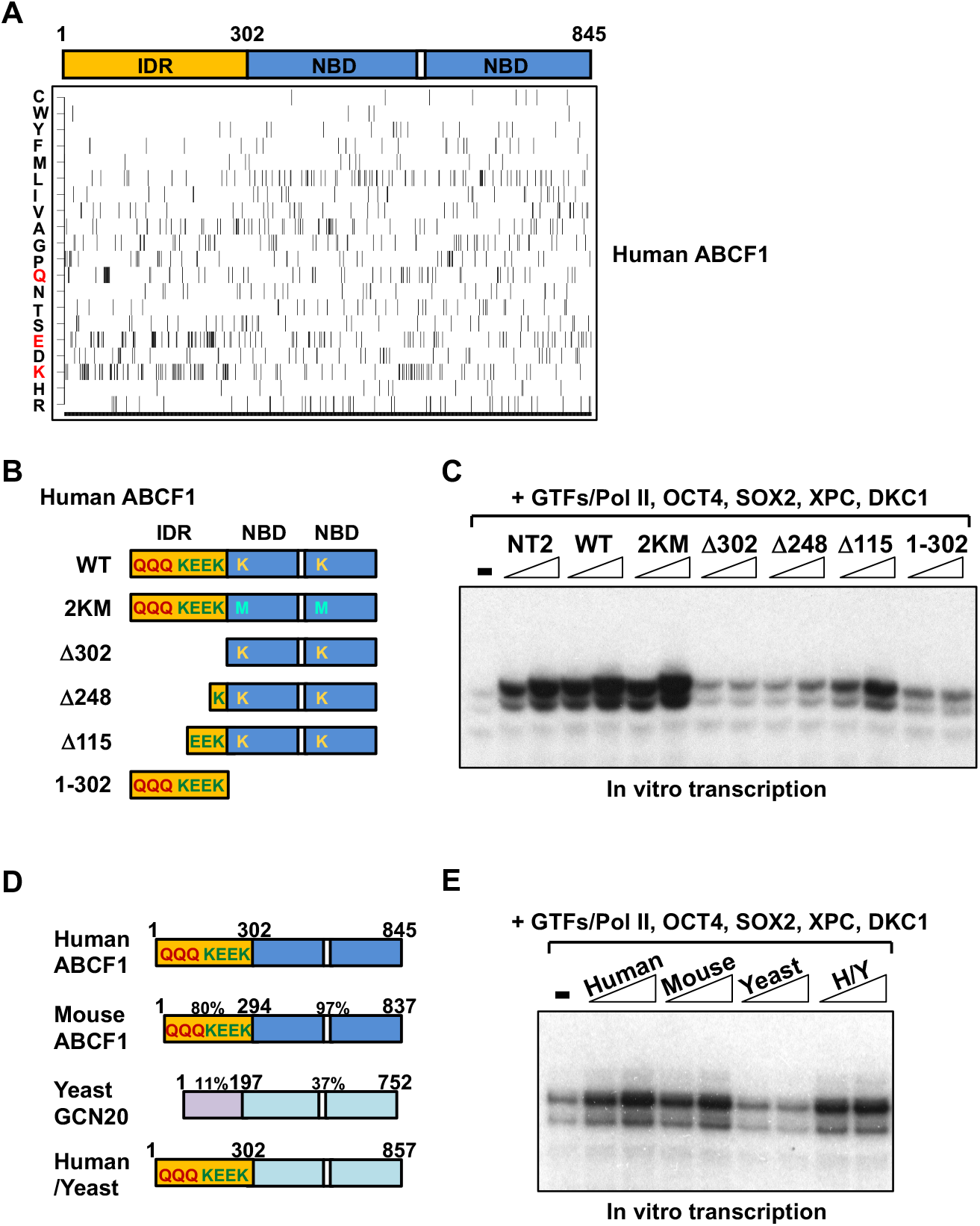
Transcriptional coactivation by ABCF1 requires its intrinsically disordered region (IDR) at the N-terminus. (A) Amino acid composition of human ABCF1. Each of the 20 amino acids is counted and marked as a black bar at that position in ABCF1. The schematic diagram (top) denotes protein domains of ABCF1: intrinsically disordered region (IDR, yellow, amino acids 1-302); two nucleotide-binding domains (NBDs, blue). One-letter abbreviations for amino acids are indicated (Left): C, Cys; W, Trp; Y, Tyr; F, Phe; M, Met; L, Leu; I, Ile; V, Val; A, Ala; G, Gly; P, Pro; Q, Gln; N, Asn; T, Thr; S, Ser; E, Glu; D, Asp; K, Lys; H, His; R, Arg. (B) Schematic diagram of full-length, wild-type (WT) ABCF1 protein depicting a predicted N-terminal IDR (yellow) containing a polyglutamine (polyQ) tract and lysine (K)/glutamic acid (E)-rich regions. The two conserved lysine residues (K324, K664) in the Walker A motif of each of the two NBDs (blue) in ABCF1 are highlighted. Full-length WT ABCF1, ATP-binding defective lysine-to-methionine mutant (2KM), various truncated ABCF1 proteins lacking part (Δ248, Δ115) or all of the IDR (Δ302) as well as the IDR by itself (1-302) are purified from *E. coli*. (C) Transcriptional activities of the various recombinant ABCF1 proteins shown in (B) are assayed (over a two-fold concentration range) together with recombinant XPC and DKC1 complexes in in vitro transcription as described in Figure 1. (D) Schematic representation of the human and mouse ABCF1, and yeast homolog GCN20. The percentage of sequence similarity among human, mouse, and yeast homolog is indicated. Domain-swapped hybrid protein between the human IDR and yeast NBDs (H/Y) is generated and purified from *E. coli*. (E) Titration over a two-fold concentration range of human and mouse ABCF1, yeast GCN20, and human-yeast hybrid (H/Y) proteins are assayed in in vitro transcription reactions. The following figure supplement is available for figure 5: Supplementary Figure S5.

The yeast homolog of ABCF1, GCN20, shares with its human and mouse counterpart a conserved NBD (Vazquez de Aldana et al., 1995). However, the residues outside of NBD (amino acids 1-197) are highly divergent from mammalian ABCF1. This domain lacks the polyQ tract and is not K/E-rich (Figures 5D and S5D). More importantly, this region is predicted to be structured, unlike the IDRs found in human and mouse ABCF1 proteins (Figure S5A). As predicted based on a requirement for the IDR, we found that purified GCN20 exhibited no coactivator activity in the in vitro transcription assay (Figures 5E and S5E). However, replacing the GCN20 N-terminal domain with the human ABCF1 IDR fully conferred transcriptional stimulatory capacity to the hybrid protein in vitro. These observations provide strong biochemical evidence that the mammalian-specific IDR confers coactivator activity.

Having identified the IDR as a “transactivation domain” in ABCF1, we next investigated the mechanisms by which the IDR potentiates stem cell-specific transcription. One of the defining features of transcriptional coactivators is their ability to act as molecular “glue” by linking transcriptional activators and other cofactors to the Pol II machinery (Boulay et al., 2017; Chan and La Thangue, 2001; Jeronimo and Robert, 2017; Sabari et al., 2018). We tested whether the IDR in ABCF1 can associate with its two co-dependent coactivators, XPC and DKC1, as well as Pol II in vitro by performing glutathione S-transferase (GST) pull-down assay. We incubated NT2 cell nuclear extract with immobilized IDR from human ABCF1 (1-302) and the transcriptionally inactive N-terminal domain from yeast GCN20 (1-197) as GST-fusion proteins. Western blot analysis revealed that only IDR/ABCF1 was able to bind XPC, DKC1, and Pol II (Figures 6A and S6A).

**Figure 6.**
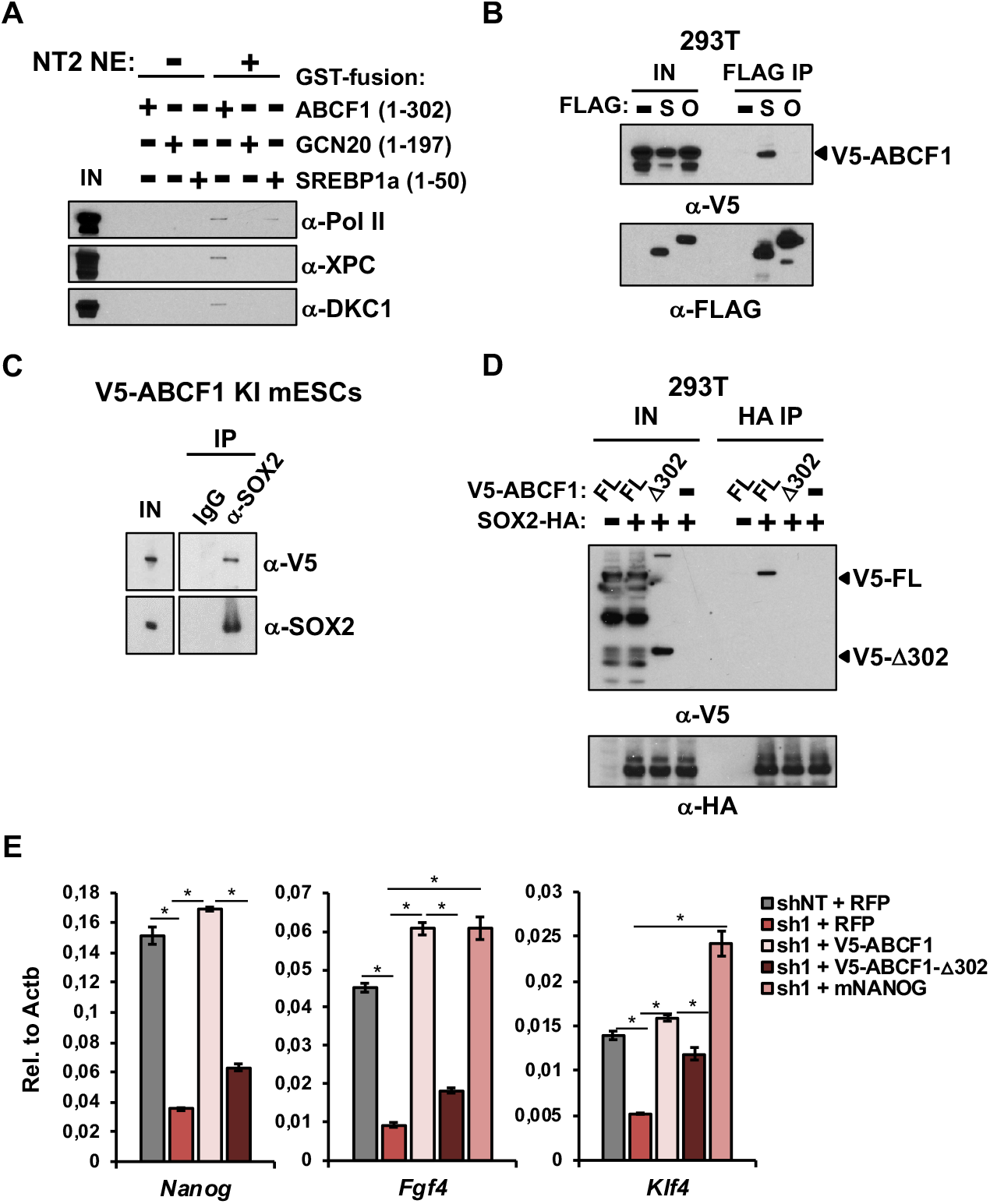
Mechanisms of transcriptional coactivation by ABCF1. (A) GST-fusion proteins containing the IDR of human ABCF1 (1-302), the N-terminal domain of yeast GCN20 (1-197), and the transactivation domain of human transcription factor SREBP1a (1-50) are incubated with buffer only (−) or NT2 nuclear extracts (NE, +). Input NE (IN) and bound proteins are analyzed by western blotting using antibodies against the largest subunit of RNA polymerase II (Pol II, RBP1), XPC, and DKC1. (B) WCEs from 293T cells co-transfected with plasmid expressing V5-ABCF1 together with either empty plasmid (−) or plasmids expressing FLAG-tagged OCT4 (O) or SOX2 (S) are immunoprecipitated with anti-FLAG antibody. Input WCEs (IN) and bound proteins are detected by western blotting. (C) Input V5-ABCF1 KI ES cell WCEs (IN) and IPs by IgG and anti-SOX2 antibodies are analyzed by western blotting using anti-V5 and anti-SOX2 antibodies. (D) HA IPs from 293T cells overexpressing HA-tagged SOX2 (SOX2-HA) with V5-tagged full-length (FL) or IDR-truncated human ABCF1 (Δ302). SOX2-bound V5-ABCF1 proteins are detected by western blotting. (E) ABCF1 knockdown-rescue assay. mRNAs from control D3 ES cells (shNT) overexpressing RFP, and ABCF1 knockdown ES cells (sh1) overexpressing RFP, V5-tagged full-length or IDR-truncated human ABCF1 (Δ302), or mouse NANOG are analyzed for *Nanog*, *Fgf4*, and *Klf4* mRNA levels by qPCR, normalized to *Actb*. Error bars present SEM. *n* = 3. (*) *P* < 0.05, calculated by two-sided Student’s t-test. The following figure supplement is available for figure 6: Supplementary Figure S6.

To further demonstrate the specificity of IDR/ABCF1 for the XPC and DKC1 complexes, we also tested a GST-fusion protein containing the transactivation domain from human transcription factor SREBP1a that is known to bind the prototypical coactivator Mediator (Näär et al., 1999).

No significant binding of XPC and DKC1 was observed, but the fusion protein was able to pull down Pol II, likely indirectly through a Mediator-Pol II interaction (Harper and Taatjes, 2018; Näär et al., 2002). Using transient transfection and co-immunoprecipitation experiments in 293T cells, we showed that ABCF1 interacted preferentially with SOX2 but not OCT4 (Figure 6B). We further confirmed the interaction between ABCF1 and SOX2 in mouse ES cell extracts (Figure 6C). Importantly, deletion of the IDR from ABCF1 (Δ302), which abrogates its transcriptional activity in vitro, completely abolished its ability to bind SOX2 (Figure 6D). Thus, the transcriptional defect observed with IDR-truncated ABCF1 is likely due to its failure to interact with SOX2, SCCs, and the Pol II machinery.

To provide in vivo evidence that the ABCF1 IDR is critical for OCT4/SOX2-dependent transcription, we performed knockdown-rescue experiments in mouse ES cells. Simultaneous knockdown of mouse ABCF1 and ectopic expression of full-length human ABCF1, but not IDR-truncated ABCF1 (Δ302) or control RFP, restored expression of several ABCF1-dependent pluripotency genes such as *Nanog*, *Fgf4*, and *Klf4* in mouse ES cells (Figures 6E and S6B). These results demonstrate that the transcriptional defect in ABCF1 knockdown ES cells is not due to off-target effects of shRNAs, and that the IDR of ABCF1 is required for pluripotency gene expression in vivo.

We next showed that ectopic expression of NANOG alone was sufficient to restore pluripotency gene expressions (*Fgf4* and *Klf4*) in ABCF1-deficient ES cells. This result indicates that *Nanog* is likely a critical downstream target of ABCF1, consistent with our in vitro transcription results. More importantly, unlike ABCF1, transcription factor NANOG has no other known roles beyond transcription in ES cells. The fact that NANOG can bypass the requirement of ABCF1 strongly suggests that the transcriptional defect observed in ABCF1 knockdown ES cells is not an indirect effect caused by disruption of other cellular pathways controlled by ABCF1. Rather, our data thus far suggest that ABCF1 directly controls pluripotency gene expression in vitro and in vivo and, by extension, stem cell maintenance. Therefore, we conclude that ABCF1 mediates stem cell-specific transcriptional activation through IDR-dependent assembly of select activator and coactivators with the Pol II transcription machinery.

### ABCF1 Links Gene Expression and DNA sensing in ES Cells

DNA sensing by the innate immune system underpins many physiological responses to DNA (Paludan and Bowie, 2013), including immunity to DNA viruses and bacteria (Chiu et al., 2009; Ishii et al., 2006; Ishikawa et al., 2009; Stetson and Medzhitov, 2006) and inflammatory responses to intracellular self-DNAs that arise from genome instability (Dunphy et al., 2018; Härtlova et al., 2015; MacKenzie et al., 2017). However, ES cells lack the ability to mount a robust pro-inflammatory response to these intracellular DNAs (D’Angelo et al., 2017; Guo et al., 2019a; Wang et al., 2013; Yu et al., 2009; Zampetaki et al., 2006). Whether there are alternative outputs in ES cells in response to aberrant intracellular DNAs remains poorly understood. Because ABCF1, an essential stem cell coactivator, also functions as an important sensor for intracellular DNAs in the innate immune system in somatic cells (Lee et al., 2013), we next asked whether ABCF1 can also recognize these DNAs in ES cells to couple DNA sensing to pluripotency gene regulation.

We took two independent approaches to test whether ABCF1 can bind intracellular DNAs in ES cells. As a first approach, we incubated mouse ES cell extracts with a 5’ biotinylated single-stranded (ss) or two different double-stranded (ds) DNA oligonucleotides containing sequences derived from *Listeria monocytogenes*. ds-matched (ds-M) contains a consensus sox-motif while ds-unmatched (ds-UM) does not. In neutrophils, SOX2 has been shown to act as a sequence-specific innate immune sensor for sox-motif-containing DNAs such as ds-M (Xia et al., 2015). Given that ABCF1 binds SOX2 in ES cells, we asked whether ABCF1 bound DNA, and if so, whether binding was direct or indirect through SOX2. Western blot analyses of protein-DNA complexes captured by streptavidin beads revealed that both ds-M and ds-UM efficiently pulled down ABCF1 while ssDNA did not (Figure 7A). These results indicate that ABCF1 binds to short dsDNAs but likely in a sequence-independent, and therefore, SOX2-independent manner. To evaluate the effect of binding of these dsDNAs by ABCF1 on pluripotency gene expression, we modeled pathogen infection in ES cells by transfecting 5’ 6-carboxyfluorescein (6-FAM)-labeled ss, ds-M, or ds-UM oligonucleotides into mouse ES cells. Transfected, 6-FAM-positive cells were enriched by flow cytometry (Figure S7A). Gene expression analyses by qPCR revealed that transfection of ds-M or ds-UM downregulated pluripotency genes and upregulated differentiation genes compared to ssDNAs (Figure S7B). These data are consistent with our observation that ABCF1 selectively binds dsDNA but not ssDNA (see Figure 7A). Importantly, the fact that both ds-M and ds-UM were equally potent in eliciting changes in pluripotency and differentiation gene expressions is distinct from what was observed in neutrophils, thus strongly suggesting a SOX2-independent mechanism. For the second approach, we investigated whether ABCF1 can bind short DNAs accumulated in cells following DNA damage-induced stress. We treated V5-ABCF1 knock-in mouse ES cells with DNA damaging agent etoposide (ETO) (Dunphy et al., 2018; Härtlova et al., 2015). Intracellular self-DNA fragments were readily detected in the cell nucleus 6 hr post-treatment (Figures 7B and S7C). We showed that immunoprecipitation of ABCF1 from ETO-treated ES cell extracts co-precipitated small DNA fragments that correspond to the size of a mononucleosome (Figure 7C). Together with in vitro data showing a direct binding of ABCF1 to naked dsDNAs, these data provide evidence that ABCF1 can also recognize short endogenous DNAs in ES cells upon DNA damage. In an effort to understand the consequence of endogenous DNA binding on the coactivator function of ABCF1, we showed that, under the same DNA damaging condition, interaction of ABCF1 with SOX2 was disrupted in ES cells (Figure 7D). These results suggest that binding of ABCF1 to self-DNA fragments accumulated in cells after DNA damage may compete with SOX2. Accordingly, we observed a loss of enrichment of ABCF1 at its target genes such as *Oct4*, *Sox2*, and *Nanog* in ETO-treated ES cells by ChIP-qPCR as early as 6 hr post ETO treatment (Figures 7E, S7D-E). Decrease in ABCF1 occupancy at promoters was accompanied by downregulation of pluripotency-associated genes and upregulation of lineage-specific genes (Figure S7F), consistent with loss of pluripotency and spontaneous differentiation in ETO-treated ES cells. If the observed trade-off in self-renewal upon DNA damage is indeed mediated through ABCF1, a prediction from this competition model is that increasing cellular concentration of ABCF1 could enhance DNA damage tolerance in ES cells, by increasing the pool of ABCF1 proteins available for transcriptional activation. Indeed, we found that ectopic expression of ABCF1, but not RFP, significantly enhanced the self-renewal capacity of ETO-treated ES cells as shown by an increase in number of AP-positive colonies formed in limiting dilution assays (Figures 7F and S7G). Importantly, we did not observe significant differences between RFP and ABCF1 gain-of-function ES cells in the absence of DNA damage, thus suggesting a specific protective effect of ABCF1 overexpression on self-renewal capacity when ES cells encounter significant genome instability. Our studies reveal a new and important safeguard mechanism, whereby the pluripotency gene network can be coupled with intracellular DNA sensing, likely by modulating the availability of ABCF1 for transcriptional coactivation.

**Figure 7.**
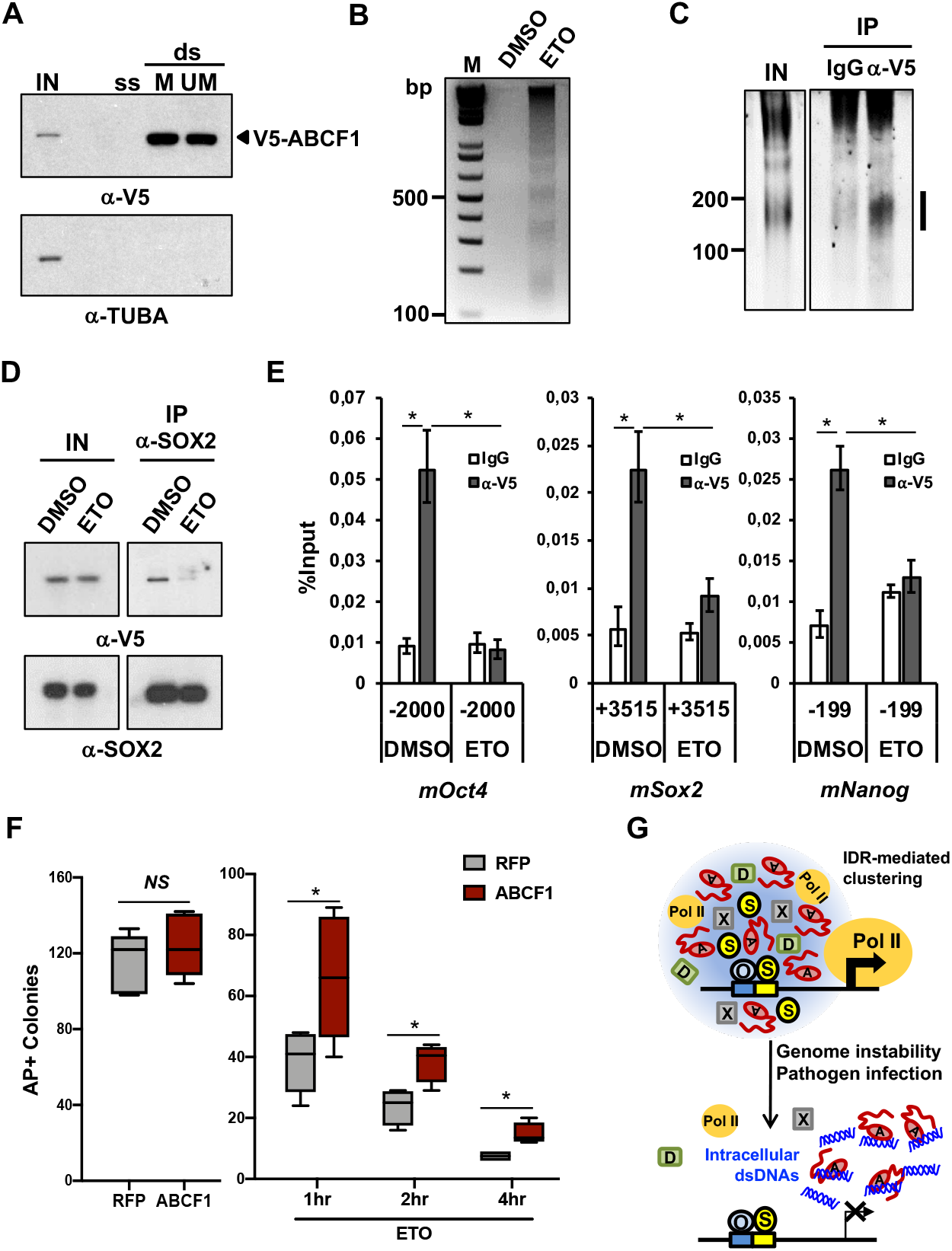
Intracellular DNAs modulate pluripotency gene expressions through ABCF1. (A) V5-ABCF1 D3 mouse ES cell WCEs are incubated with three different 5’ biotinylated 98mer oligonucleotides: single-stranded (ss), double-stranded (ds) with SOX2-binding motif (Matched, ds-M), or ds without the motif (Unmatched, ds-UM). These DNA sequences are derived from *Listeria monocytogenes* genome. Input WCEs (IN) and streptavidin-beads captured, DNA-bound ABCF1 proteins are analyzed by western blotting. α-tubulin (TUBA) is used as control for binding specificity. (B) Genomic DNA purified from nuclear extracts prepared from DMSO and etoposide-treated (ETO, 20 μM) V5-ABCF1 knock-in (KI) D3 ES cells were analyzed on agarose gel and stained with ethidium bromide. (C) WCEs prepared from ETO-treated (20 μM) V5-ABCF1 KI D3 ES cells are incubated with IgGs or anti-V5 antibodies. Co-purified nucleic acids are treated with RNase A, separated on urea-PAGE, and stained with SYBR Gold. Vertical bar denotes DNAs specifically bound by ABCF1. (D) DNA damage disrupts ABCF1-SOX2 interaction. Input (IN) and SOX2 IPs from WCEs of DMSO or ETO-treated (20 μM) V5-ABCF1 KI D3 ES cells are analyzed by western blotting. (E) MNase-ChIP of ABCF1 in DMSO and ETO-treated (80 μM) V5-ABCF1 KI D3 ES cells. Enrichment of ABCF1 on OCT4/SOX2-targeted regions of *Oct4*, *Sox2*, and *Nanog* gene promoters is analyzed by qPCR as in Figure 4. (F) Colony formation assays in control and ABCF1 gain-of-function D3 cells. 200 D3 ES cells stably expressing RFP or V5-ABCF1 are plated on 24-well plates, treated with DMSO (left) or ETO (1 μM, right) for indicated period of time (hr), and let recover for 6 days before staining for AP activity. AP-positive colonies are counted. Error bars represent SEM of three independent experiments. *n* = 3. (*) *P* < 0.05, calculated by two-sided Student’s t-test. (G) Model depicting mechanisms whereby ABCF1 couples pluripotency gene transcription with intracellular DNA sensing. ABCF1 IDR (red wavy line) promotes specific clustering and formation of a hub comprising of SOX2 (S), XPC (X), DKC1 (D), and Pol II molecules at target gene promoter to stimulate transcription by increasing local concentration of these factors. ABCF1 proteins available for transcription are diverted to bind intracellular dsDNAs generated from genome instability or pathogen infection. Decrease in ABCF1 at gene promoters destabilizes the multivalent interactions between SOX2, XPC, DKC1, and Pol II. This leads to disruption of the protein hub and decrease in gene transcription by Pol II. Downregulation of pluripotency-associated genes promotes differentiation of compromised ES cells and their elimination from the self-renewing population. The following figure supplement is available for figure 7: Supplementary Figure S7.

## DISCUSSION

### Role of IDR in Mediating Stem Cell-specific Transcriptional Activation

In this study, we identified ABCF1 as a critical transcriptional coactivator for stem cell maintenance. ABCF1 plays a pivotal role in linking co-dependent SCCs (XPC and DKC1) with SOX2 and the Pol II machinery, thus providing a mechanistic basis for SCC-dependent transcriptional activation by OCT4 and SOX2. The three SCCs are not expressed exclusively in ES cells. This raises an important question of why ES cells employ these coactivators, which are also expressed in somatic cells, to drive stem cell-specific transcriptional activation and the stem cell fate. Our studies reveal several unique properties of SCCs that may contribute to their gene-specificity (Fong et al., 2011, 2014). All three SCCs are highly enriched in ES cells. In addition, we demonstrated that SCCs synergistically stimulate transcription in vitro, where omission of ABCF1 from in vitro transcription reactions significantly decreased transcriptional coactivation by XPC and DKC1 (Figures 1B and 5C). This is supported by our in vivo observations that knockdown of ABCF1 resulted in downregulation of a number of key pluripotency genes (Figure 3C). Most importantly, SCCs appear to be able to physically and functionally interact with each other and with select stem cell-specific transcription factors such as OCT4 and SOX2, where ABCF1 likely plays an important role in mediating many of these interactions (Figure 6). For example, we showed that the IDR in ABCF1 is critically important for both physical interaction with SOX2, XPC, DKC1, and Pol II, and functional reconstitution of ABCF1 coactivator activity both in vitro and in vivo (Figures 5E and 6E). The flexible nature of IDRs is thought to facilitate the dynamic interaction with multiple protein partners, by virtue of their ability to rapidly adopt an ensemble of conformations (Choi et al., 2019; Zhang and Tjian, 2017). It is worth emphasizing that IDRs are not simply unstructured sequences that bind promiscuously to any proteins; instead, they can be selective for binding partners (Chong et al., 2018; Guo et al., 2019b). In this regard, the conformationally flexible XPC protein also contains several highly disordered regions that we found, however, to be dispensable for transcriptional activation (Cattoglio et al., 2015; Fong et al., 2011; Zhang et al., 2015) (data not shown). These observations reveal the unique ability of ABCF1 IDR to integrate multiple lines of information encoded by SOX2, SCCs, and the Pol II machinery, likely by forming a hub of these factors at target gene promoters through selective multivalent interactions (Figure 7G) (Chong et al., 2018). This is analogous to previous reports suggesting that IDRs found in other coactivators may employ a protein:protein driven local high concentration mechanism to activate transcription (Ann Boija et al., 2018; Boulay et al., 2017; Chong et al., 2018; Sabari et al., 2018). Our findings here also provide a clear example of an IDR that can impart activator preference and target gene specificity to a transcriptional coactivator.

### An Alternative Mechanism for ES Cells to Sense Infection and Genome Instability

Studies have shown that ES cells may be able to combat infections without a robust innate immune system by constitutively expressing a subset of interferon-stimulated genes (ISGs) that function to suppress certain viral infections. However, this handful of ISGs are likely effective against only a subset of pathogens (Wu et al., 2018). High proliferation rate of mouse ES cells and a short gap (G1) phase of the cell cycle impose replication stress that, in turn, can lead to genome instability and DNA damage (Ahuja et al., 2016; Aladjem et al., 1998; Suvorova et al., 2016). While ES cells do express other known DNA sensors (e.g. cGAS and STING, data not shown) that in theory can detect intracellular DNAs from infection and DNA damage, their downstream signaling pathways required to activate a robust pro-inflammatory response are absent or highly attenuated, in part due to active suppression by stem cell transcription factors including OCT4 and SOX2 (Eggenberger et al., 2019; Guo, 2019). Nonetheless, studies have indicated that ES cells are hypersensitive to DNA damage and readily undergo differentiation and apoptosis (Aladjem et al., 1998; Heyer et al., 2000). This is not due to a diminished DNA repair capacity because many repair-related factors are in fact elevated in ES cells (Choi et al., 2017; Maynard et al., 2008; Vitale et al., 2017). Taken together, these observations suggest that alternative mechanisms likely exist in order to sensitize ES cells to these cellular insults, thereby protecting the pristine state of ES cells from infections and genome instability.

### Biological Implications of ABCF1/IDR-mediated Pluripotency Gene Expression

Because IDR-mediated interactions are highly dynamic (Chong et al., 2018; Hnisz et al., 2017; Staby et al., 2017; Weng and Wang, 2020; Zhang and Tjian, 2017), targeting these transient and multivalent interactions between transcription factors and SCCs could provide an effective means of modulating the pluripotency gene network in a rapid manner in response to changing cellular signals. Indeed, it has been shown that molecular crowding by IDR is particularly sensitive to changes in concentration, where a less than two-fold decrease can be sufficient to cause an IDR-mediated phase-separated body to be disrupted (Wang et al., 2018). Therefore, we suggest that a decrease in ABCF1 protein concentration, such as during stem cell differentiation (Figure 2) or sequestration from SOX2 by intracellular DNAs during DNA damage or infection (Figure 7), could lead to rapid dissolution of the transcriptional apparatus, pluripotency gene expression, and the pluripotent state as a result. This rapid transcriptional response is likely a necessary feature of ES cells where wholesale changes in gene expression underlie commitment to exit of pluripotency and cellular differentiation during development. Likewise, in the event that DNA damage or infections exceeds the capacity of ES cells to repair or clear, compromised cells can be eliminated from the self-renewing pool through spontaneous differentiation.

How are IDR-mediated interactions of SCCs regulated? For example, we observed that binding of ABCF1 with SOX2 and to SOX2-target gene promoter remained largely unaffected under lower ETO concentrations. Accordingly, expression of core pluripotency genes such as *Oct4*, *Sox2*, *Nanog* were not significantly downregulated. Only when a relatively high concentration of ETO or oligonucleotides was used did we observe significant disruption in binding and gene expression (data not shown). Based on these observations, we suggest that this sigmoidal-like response of pluripotency gene expression to DNA damage and intracellular DNA levels can be explained at least in part by ABCF1-SOX2 binding behavior. Under optimal growth conditions, there are multiple mechanisms to return ES cells experiencing low level of DNA damage or infections to homeostasis, such as DNA repair or constitutive ISG expressions. We surmise that when these cellular insults turn catastrophic, ABCF1 serves as a critical checkpoint for self-renewal to ensure that compromised ES cells do not propagate, by enforcing their exit of pluripotency and into differentiation. This safeguard mechanism allows ES cells to robustly stabilize the pluripotency gene network while remaining responsive to pathogen infections and genome instability.

We should emphasize that the model proposed here entails that ABCF1 is limiting in ES cells (Figure 7G). This is likely the case because ectopic expression of ABCF1 significantly improved the self-renewal capacity of ES cells under DNA damaging condition but had no discernible beneficial effects in undamaged cells. These observations place ABCF1 directly in this crosstalk and are consistent with our hypothesis that ABCF1 becomes limiting for transcriptional coactivation when challenged with endogenous DNA fragments induced by DNA damage (Figure 7G). Based on our findings, we propose that ABCF1 represents an important regulatory nexus, wherein the constant tug-of-war between transcriptional activation and intracellular DNA sensing by ABCF1 could drive an ES cell to self-renew or commit to differentiation. This switching of cell fates critically depends on whether intracellular DNA rises above a certain threshold level that irreversibly tilts the balance toward rapid exit of pluripotency.

## MATERIALS AND METHODS

### DNA constructs and antibodies

cDNAs for human and mouse ABCF1 were obtained from cDNA libraries generated from total RNAs isolated from human embryonic carcinoma cell line NTERA-2 (NT2) and mouse embryonic stem (ES) cell line D3. Yeast GCN20 cDNA was amplified from purified S. cerevisiae genomic DNA. For expression of GCN20, full-length human and mouse ABCF1, as well as various truncations of human ABCF1 in *E. coli*, corresponding cDNAs were cloned into a modified pMtac-His6 vector containing a His6 tag at the N-terminus and a FLAG tag at the C-terminus. Mutations of the two Walker-A motifs in human ABCF1 (K324M, K664M) were created by using Quikchange II Site Directed Mutagenesis Kit (Agilent). Human ABCF1 and yeast GCN20 domain fusion cDNAs were created by PCR-mediated ligation and cloned into modified pMtac-His6 vector. GST fusion proteins containing N-terminal domain of human ABCF1 (amino acids (aa) 1-302), yeast GCN20 (aa 1-197), and transcription factor SREBP1a (aa 1-50) were generated by inserting the corresponding cDNA fragments into pGEX4T-3 vector (Sigma). V5-tagged full-length and N-terminal truncation (Δ302) of human ABCF1, and untagged mouse *Nanog* were cloned into lentiviral overexpression vector pHAGE-IRES-Neo (Fong et al., 2011). cDNAs for human OCT4 and SOX2 were cloned into the pFLAG-CMV5a mammalian expression vector (Sigma).

Polyclonal antibodies against ABCF1 (13950-1-AP) was purchased from ProteinTech, XPC (A301-122A) and mouse Nanog (A300-397A) from Bethyl Laboratories, DKC1 (H-300), OCT4 (N-19), and Pol II (N-20) from Santa Cruz Biotechnology, SOX2 (AB5603) from EMD Millipore, V5 ChIP grade (ab15828) from Abcam. Monoclonal antibodies against ACTB (66009-1) from ProteinTech, HA-tag (C29F4) from Cell Signaling Technology, FLAG-tag (M2) and α-Tubulin (T5168) from Sigma, RFP (600-401-379) from Rockland, and V5-tag (R96025) from Life Technologies.

### Cell culture

NT2 cell line was obtained from ATCC. NT2, 293T, and HeLa cells were cultured in DMEM high glucose with GlutaMAX (Gibco) supplemented with 10% fetal bovine serum (FBS). Large scale culture of NT2 cells were described (Fong et al., 2011). Feeder-independent human ES cell line H9 was purchased from WiCell Research Institute. H9 ES cells were cultured in mTeSR1 (STEMCELL Technologies) with normocin (50 *μ*g/ml; Invivogen) on Matrigel-coated tissue culture plates (Corning). D3 mouse ES cell line was purchased from ATCC and adapted to feeder-free condition as described (Fong et al., 2011). Medium was changed daily. Cell cultures were passaged by StemPro accutase (Gibco) for human ES cells and trypsin for mouse ES cells.

For chromatin immunoprecipitation (ChIP) experiments, mouse ES cells were adapted to 2i/LIF culture condition. Mouse ES cells were passaged in 2i/LIF, serum-free medium composed of 1:1 mix of Advanced-DMEM /F12 and Neurobasal medium (Gibco) supplemented with N2 (Gibco), B27 (Gibco), L-glutamine (GlutaMAX, Gibco), beta-mercaptoethanol (0.1 mM; Sigma), BSA (50 *μ*g/ml; Sigma), PD0325901 (1 *μ*M; Sigma), CHIR99021 (3 *μ*M; EMD Millipore), and LIF (10^2^ U/ml) for at least four passages before the cells were used for ChIP and RT-qPCR analyses. Differentiation of H9 ES cells was induced by exchanging human ES cell growth medium with DMEM/F12 (Gibco) containing 2 mM L-glutamine, 12.5% FBS, and normocin (50 *μ*g/ml) for up to 14 days. V5-ABCF1 knock-in D3 mouse ES cells were induced to differentiate by maintaining cells in regular medium containing 5 *μ*M all-trans retinoic acid (Sigma) for 7 days.

### In vitro transcription assay

In vitro transcription reactions, the human *NANOG* transcription template, purification of activators OCT4 and SOX2, general transcription factors, RNA polymerase II, and recombinant XPC complex were described (Fong et al., 2011). Recombinant XPC complex purified from Sf9 and DKC1 complex reconstituted and purified from Sf9 cells or *E. coli* (Fong et al., 2014) were supplemented in the in vitro transcription reactions to assay for SCC-B activity.

### Purification of SCC-B/ABCF1

All steps were performed at 4°C. Nuclear extracts were prepared from 400 L of NT2 cells. Partially purified P11-phosphocellulose 1 M KCl and Ni-NTA flowthrough (Ni-FT) fractions were prepared as described (Fong et al., 2011). The Ni-FT fraction was dialyzed against buffer D (20 mM HEPES, pH 7.9, 2 mM MgCl_2_) at 0.2 M KCl with 0.0025% NP-40 and 10% glycerol (all buffers from then on contained 0.0025% NP-40 and 10% glycerol unless otherwise stated). This Ni-FT fraction was applied to a Poros 20 HQ column (Applied Biosystems), subjected to a 4 column volume (CV) linear gradient from 0.2 M to 0.4 M KCl (Q0.3), washed at 0.52 M KCl, and developed with a 13 CV linear gradient from 0.52 M to 1.0 M KCl. Transcriptionally active Q0.3 fraction (0.32–0.4 M) were pooled and applied directly to hydroxyapatite (HAP) type II ceramic resin (Bio-Rad), washed first at 0.38 M, then lowered to 0.1 M KCl in 3 CV. HAP column buffer was then exchanged and washed extensively with buffer D at 0.03 M KPi, pH 6.8 without KCl and NP-40. The HAP column was subjected to a 20 CV linear gradient from 0.03 M to 0.6 M KPi. Active HAP fractions eluting from 0.2–0.3 M KPi were pooled and separated on a Superose 6 XK 16/70 gel filtration column (130 ml, GE Healthcare) equilibrated with buffer D + 0.1 mM EDTA at 0.15 M KCl. Active Superose 6 fractions with an apparent molecular mass of 400 - 600 kDa were pooled and supplemented with 0.25 mg/ml insulin (Roche). Pooled fractions were applied to a Poros 20 HE column (Applied Biosystems) equilibrated in buffer D + 0.1 mM EDTA at 0.15 M KCl, subjected to a 34 CV linear gradient from 0.15 M to 1 M KCl. SCC-B containing HE fractions eluted from 0.35 - 0.43 M KCl. Active HE fractions eluting from 0.35-0.43 M KCl were supplemented with 0.3 mg/ml insulin and dialyzed to 0.15 M KCl in buffer D + 0.1 mM EDTA + 0.01% NP-40. The dialyzed HE fraction was applied to a Mono S PC 1.6/5 column (GE Healthcare), washed and developed with a 20 column volume linear gradient from 0.15 M to 0.65 M KCl. Transcriptionally active SCC-B/ABCF1 fractions eluted from 0.29 – 0.31 M KCl.

### Mass spectrometry analysis

Peak Mono S fractions were pooled, concentrated using a Spin-X centrifugal concentrator, separated by SDS-PAGE, stained with PageBlue (Thermo Fisher), protein bands excised, digested with trypsin, and extracted. Peptide pools from each gel slice were analyzed by matrix-assisted laser desorption time-of-flight mass spectrometry (MALDI-TOF MS; Bruker Reflex III). Selected mass values were used to search protein databases linked to PROWL (Rockefeller University) using ProFound and protein databases linked to ExPASy (Swiss Institute of Bioinformatics, Geneva) using PeptIdent.

### Generation of endogenously V5-tagged ABCF1 knock-in mouse ES cell line

sgRNA targeting genomic region immediately downstream of the ATG translation start codon of ABCF1 was cloned into LentiCRISPRv2 vector (Addgene). A single-stranded (ss) donor oligonucleotide containing a V5-tag followed by a flexible linker GSSG sequence in frame with the second amino acid of ABCF1, which is flanked by left and right homology arms of about 70 bp, was synthesized (Integrated DNA Technologies (IDT)). Both the LentiCRISPRv2-sgRNA plasmid and the ss donor oligonucleotides were transfected into D3 mouse ES cell line using lipofectamine 3000 (Invitrogen). Transfected cells were selected with puromycin (1.5 *μ*g/ml) for 3 days. Cells were then expanded in the absence of puromycin to avoid integration of the LentiCRISPRv2-sgRNA plasmid into the genome. Single clones were plated into 96-well plates. Positive clones were identified by PCR and confirmed by sequencing and western blotting. Clones selected for further analysis were confirmed to be puromycin-sensitive and express similar levels of key pluripotency genes such as *Nanog*, *Oct4*, and *Sox2* (Figure S4B). See Table S1 for sgRNA and ss donor oligo sequences.

### Purification of recombinant proteins

pMtac expression plasmids were transformed into BL21-Codon Plus RIPL competent cells (Agilent). Expression of His6-tagged proteins were induced at 16°C overnight with 0.25 mM IPTG. Cell pellets were lysed in high salt lysis buffer HSLB (50 mM Tris–HCl, pH 7.9, 0.5 M NaCl, 0.6% Triton X-100, 0.05% NP-40, 10% glycerol) with imidazole (10 mM) and lysozyme (0.5 mg/ml). Sonicated lysates were cleared by ultracentrifugation and incubated with Ni-NTA resin for 16 hr at 4°C. Bound proteins were washed extensively with HSLB with 20 mM imidazole, equilibrated with 0.3 M NaCl HGN (25 mM HEPES, pH 7.9, 10% glycerol, 0.05% NP-40) with 20 mM imidazole, and eluted with 0.25 M imidazole in 0.3 M NaCl HGN. Peak fractions were pooled and incubated with anti-FLAG M2 agarose (Sigma) for 3 hr at 4°C. Immunoprecipitated proteins were washed extensively with 0.7 M NaCl HEMG (25 mM HEPES, pH 7.9, 0.1 mM EDTA, 12.5 mM MgCl_2_,10% glycerol) with 0.1% NP-40 and equilibrated with 0.25 M NaCl HEMG with 0.1% NP-40. Bound proteins were eluted in the same buffer containing 0.4 mg/ml FLAG peptides. Eluted proteins were filtered through a 0.22 *μ*m filter, snap frozen in liquid nitrogen, and stored at −80°C.

pGEX4T-3 expression plasmids were transformed into BL21-Codon Plus RIPL competent cells (Agilent). Protein expression was induced at 20°C for 3.5 hr. Bacterial pellets were resuspended in 1X PBS containing 3 M DTT, 0.5 mM PMSF, and complete protease inhibitor cocktail (Sigma) and snap frozen in liquid nitrogen. Sonicated lysates were cleared by ultracentrifugation. Cleared lysates were then incubated with Glutathione Sepharose 4 Fast Flow (GE Healthcare) to immobilize GST-fusion proteins.

### GST pull-down assay

Nuclear extracts from NT2 cells were prepared as described (Dignam et al., 1983). Proteins were precipitated with ammonium sulfate (55% saturation) and resuspended in buffer D (20 mM HEPES, pH 7.9, 2 mM MgCl_2_, 20% glycerol) containing 20 mM KCl and 0.01% NP-40 to about a fourth of the starting volume of nuclear extracts. Soluble extracts were dialyzed against buffer D containing 0.2 M KCl, 0.2 mM EDTA, 10% glycerol, and 0.01% NP-40 (buffer HEGN). Dialyzed nuclear extracts were cleared by centrifugation.

Bacterial lysates containing GST-fusion proteins were immobilized onto Glutathione Sepharose 4 Fast Flow in 1X PBS and 30 *μ*g/ml BSA. Bound proteins were washed extensively with STE buffer (20 mM Tris pH 8.0, 1 mM EDTA, 1% TX-100, 0.5% NP-40, 10% glycerol) containing 1M NaCl. Approximately 20 *μ*g of GST-ABCF1 (1-302), equimolar of GST-GCN20 (1-197), and GST-SREBP1a (1-50) were immobilized onto the sepharose beads. Bound GST-fusion proteins were equilibrated to 0.2 M NaCl HEGN.

Approximately 3 mg of NT2 nuclear extracts were incubated with the immobilized GST-fusion proteins overnight at 4°C. Bound proteins were washed extensively with 0.3 M NaCl HEGN, then with 0.1M NaCl HEGN. Proteins bound to GST-fusion proteins were eluted by incubating the sepharose slurry twice with 0.1 M NaCl HEGN containing 0.2% sarkosyl.

### Coimmunoprecipitation assay

pHAGE-V5-ABCF1 (full-length and Δ302), pFLAG-CMV5a-OCT4, and pFLAG-CMV5a-SOX2 expression plasmids were co-transfected into 293T cells using Lipofectamine 2000 (Invitrogen). Transfected cells on 10 cm dishes were lysed with 1 ml of lysis buffer (0.25 M NaCl, 20 mM HEPES, pH 7.9, 2 mM MgCl_2_, 0.5% NP-40, 10% glycerol) 40 hr post transfection. Cell lysates were collected and homogenized by passing through a 25-gauge needle 5 times. Lysates were cleared by centrifugation at 15k rpm for 25 min at 4°C. Cleared lysates were then incubated with anti-FLAG M2 agarose pre-blocked with 5 mg/ml BSA for 3-4 hr at 4°C. Bound proteins were washed extensively with lysis buffer followed by FLAG peptide elution (0.4 mg/ml) in buffer HEMG containing 0.3 M NaCl and 0.1% NP-40.

To detect interaction between endogenous ABCF1 and SOX2 under normal and DNA damage conditions, V5-ABCF1 knock-in D3 mouse ES cells were treated with DMSO or etoposide (ETO, 20 *μ*M; Sigma) for 6 hr. Whole cell extracts were prepared using lysis buffer (20mM HEPES, pH 7.9, 0.12 M NaCl, 1 mM EDTA, 1% NP-40, 1 mM DTT, 1 mM Benzamidine, 0.5 mM PMSF, complete protease inhibitors (Sigma)). Cell lysates were incubated on ice for 10 min and lysates were cleared by centrifugation. Cleared lysates were incubated with anti-SOX2 antibodies at 4°C overnight. Lysates were incubated with Protein A Sepharose (GE Healthcare) for 1-2 hr at 4°C and bound proteins were washed extensively with lysis buffer and eluted with SDS sample buffer with boiling.

### Chromatin immunoprecipitation (ChIP)

V5-ABCF1 knock-in D3 mouse ES cells were first adapted to 2i/LIF condition. Conditions for crosslinking cells first with ethylene glycol bis[succinimidylsuccinate] (EGS, 3 mM; Pierce) and then formaldehyde, sonication of chromatin, and immunoprecipitation with rabbit IgG or polyclonal anti-V5 antibody (Abcam) were described (Fong et al., 2014). Input chromatin and immunoprecipitated DNA were reversed crosslinked at 65°C overnight. DNA was purified by using a Qiagen PCR purification kit (Qiagen) or by phenol/chloroform extraction followed by chloroform extraction. DNA was precipitated by adding 1/10 volume of 3M sodium acetate, pH 5.5 and 3 volumes of ethanol. Precipitated DNA was washed once with 70% ethanol and resuspended in 1X TE buffer (10 mM Tris-HCl, pH 8.0, 0.1 mM EDTA). Purified DNA was quantified by real time PCR with KAPA SYBR FAST qPCR Master Mix (KAPA Biosystems) and gene specific primers using a CFX Touch Real-Time PCR Detection System (Bio-Rad). Primer sequences are shown in Table S2.

### Micrococcal Nuclease (MNase)-ChIP

V5-ABCF1 knock-in D3 mouse ES cells were first adapted to 2i/LIF conditions. Cells were crosslinked with EGS for 20 min and then with formaldehyde for 5 min. Nuclei were prepared as described (Fong et al., 2014). Nuclei were equilibrated with MNase buffer (10 mM Tris-HCl, pH 7.5, 15 mM NaCl, 60 mM KCl, 1mM CaCl_2_, 1 mM PMSF, complete protease inhibitors (Sigma)) before cells were digested with MNase in the same buffer as described (Mieczkowski et al., 2016). Briefly, nuclei from 4 x 10^7^ of cells were resuspended in 0.5 ml of MNase digestion buffer and digested with 2400 U of MNase (NEB) in the presence of 3.3 mM CaCl_2_ and 0.2% Triton X-100 at 30°C for 12 min with vigorous shaking (1200 rpm). Digestion was terminated by adding 12.5 *μ*l of MNase stop buffer (250 mM EDTA, 250 mM EGTA, 0.5% SDS, 125 mM NaCl). MNase-digested nuclei slurry was sonicated using Bioruptor 300 (Diagenode) to disrupt the nuclear membrane. Released chromatin was clarified by centrifugation. ChIP was conducted as described above. Primer sequences are shown in Table S2.

### shRNA-mediated knockdown and rescue by lentiviral infection

For lentivirus production, non-targeting control and pLKO plasmids targeting mouse ABCF1 (Sigma), or pHAGE plasmids for overexpression were co-transfected with packaging vectors (psPAX2 and pMD2.G, Addgene) into 293T cells using lipofectamine 2000 (Invitrogen). Supernatants were collected at 48 hr and again at 72 hr. Virus preparation was performed as described (Fong et al., 2011). Functional viral titer was determined by transduction of limited dilution of concentrated viruses into HeLa or NIH 3T3 cells followed by counting antibiotic-resistant colonies to obtain colony formation unit (CFU)/ml. Cells were infected with viruses in the presence of 8 *μ*g/ml polybrene. For knockdown experiments in mouse ES cells, lentiviruses expressing control non-targeting shRNA (NT), or two independent shRNAs targeting ABCF1 were used to infect cells. Infected cells were selected with puromycin (1.5 *μ*g/ml). Pluripotency status of control and ABCF1-knockdown cells were analyzed using an alkaline phosphatase (AP) detection kit (EMD Millipore), or by RT-qPCR analysis of mRNAs purified using TRIzol reagent (Life Technologies).

For rescue experiments in ABCF1-knockdown mouse ES cells, mouse ES cells were co-infected with viruses expressing shABCF1 (pLKO-shABCF1-1) at multiplicity of infection (MOI) of 15 and viruses expressing either control RFP at MOI of 6, mouse Nanog at MOI of 6, full-length hABCF1 (FL), or N-terminally truncated ABCF1 (Δ302) (pHAGE-V5-ABCF1) at MOI of 6 and 1, respectively. We used a lower MOI for Δ302 because it expresses at a substantially higher level than full-length ABCF1 in mouse ES cells even at MOI of 1 (see supplementary figure S6B). Co-Infected cells were selected with puromycin (1 *μ*g/ml) and neomycin (750 *μ*g/ml).

### siRNA-mediated knockdown in human ES cells

For knockdown of ABCF1 in human ES cells, H9 cells were transfected with 3.3 *μ*M of si-non-targeting (siNT) and siABCF1 (siGENOME SMARTpool, Dharmacon) using Lipofectamine 3000 (Invitrogen) according to manufacturer’s instructions. 48 hr after transfection, cell viability and morphology were documented before RNAs were collected by TRIzol reagent (Life Technologies) and processed for RT-qPCR analysis.

### Somatic cell reprogramming and flow cytometry

CF-1 MEFs (Charles River) were transduced with inducible S TEMCCA and rtTA lentivirus-containing supernatants overnight in 8 μg/ml polybrene (Sigma). Doxycycline (2μg/ ml; Sigma) was supplemented to complete mouse ES cell media to induce expression of OCT4, KLF4, SOX2, and c-M YC. Reprogramming was assayed by AP staining (EMD Millipore), or by flow cytometry analysis using anti-SSEA1 (Biol egends) on a BD LSRFortessa, performed according to the manufacturers’ protocols.

### RNA isolation, reverse transcription and real time PCR analysis

Cells were rinsed once with 1X PBS. Total RNA was extracted and purified using TRIzol reagent (Life Technologies) followed by DNase I treatment (Invitrogen). cDNA synthesis was performed with 1 *μ*g of total RNA using iScript cDNA Synthesis Kit (Bio-Rad) and diluted 10-fold. Real time PCR analysis was carried out with iTaq UniverSYBR green (Bio-Rad) and gene specific primers using the CFX Touch Real-Time PCR Detection System (Bio-Rad). Results were normalized to *β-actin*. Primer sequences are shown in Table S3.

### Nucleofection of oligonucleotides and flow cytometry

D3 mouse ES cells were detached and washed once with 1X PBS. For each nucleofection 3 x 10^6^ cells were resuspended in 80 *μ*l of mouse ES cell nucleofection solution (Lonza). Equimolar (0.32 nmol) of 5’ 6-carboxyfluorescein (6-FAM) labeled ss, ds-M, or ds-UM DNA oligos (IDT) (Table S4) were nucleofected into mouse ES cells using an Amaxa nucleofector (Lonza) according to manufacturer’s instructions. Cells mock-nucleofected with water were used as a negative control. Nucleofected cells were recovered in 1 ml of pre-warmed ES cell medium at 37°C for 10 min before cells were transferred to 10 cm culture plates. 24 hr after plating, cells were detached, rinsed with 1X PBS, resuspended in 0.2 ml of ice-cold PBS, and filtered into BD filter cap tubes for fluorescent activated cell sorting (FACS). FITC-positive cells were sorted on a BD FACS Aria II instrument using the following settings: 70*μ*M nozzle, 70psi pressure, frequency 85, amplitude 5, and 70 sheath pressure. Mock-nucleofected negative control cells were used for setting up the gate for FITC+ cells. Once the gate is adjusted, FITC+ cells were collected directly into TRIzol (Life Technologies) solution and immediately frozen in dry ice.

### Cytoplasmic and nuclear fractionation

DMSO or ETO treated mouse ES cells were washed with PBS twice and lysed with Buffer A (10 mM HEPES, pH7.9, 10 mM KCl, 0.1 mM EDTA, 0.4% NP-40, 1 mM DTT, complete protease inhibitors (Sigma)) on ice for 10min. Lysates were cleared by spinning down at 15k g for 3 min at 4°C. Supernatants were kept as cytoplasmic fractions. Nuclei were washed with Buffer A 3 times and lysed with Buffer B (20mM HEPES, pH7.9, 0.4 M NaCl, 1 mM EDTA, 10% Glycerol, 1 mM DTT, complete protease inhibitors (Sigma)) with vigorous shaking for 2 hr or overnight at 4°C. Lysates were cleared by centrifugation at 15k g for 5min at 4°C. Supernatants were kept as nuclear fractions.

### ABCF1-DNA interaction analysis

For DNA pull-down assay, approximately 5 x 10^6^ V5-ABCF1 knock-in D3 mouse ES cells were lysed in 0.2 ml of lysis buffer (0.14 M NaCl, 50 mM HEPES, pH 7.9, 1 mM EDTA, 2 mM MgCl_2_, 0.5% NP-40, 10% glycerol). Cell lysates were homogenized by passing through a 25-gauge needle 5 times. Cell lysates were cleared by centrifugation. 5’ biotinylated single stranded (ss) or doubled stranded (ds) oligonucleotides labeled with biotin at the 5’ end of the sense strand was synthesized. All oligos contain modified, exonuclease-resistant nucleotides (phosphorothioate bonds) in the last 5 nucleotides at the 5’ and 3’ ends of both sense and antisense strands (IDT). These oligonucleotides were incubated with cell lysates for 1.5 hr at 4°C. For each pull down reaction, 100 *μ*l of Dynabeads MyOne Streptavidin T1 bead slurry was first pre-blocked in lysis buffer containing 50 mg/ml BSA for 1 hr at 4°C. The pre-blocked beads were then incubated with cell lysate containing oligonucleotides for 1 hr at 4°C. Biotinylated DNA oligonucleotides and associated protein factors captured onto the beads were washed extensively with lysis buffer. Bound proteins were eluted by boiling the beads in sample buffer. Sequences of the 5’ biotinylated oligonucleotides are available in Table S4.

To analyze binding of ABCF1 to endogenous dsDNA fragments, V5-ABCF1 knock-in D3 mouse ES cells were treated with DMSO or ETO (20 *μ*M) for 6 hr to induce DNA fragmentation. Whole cell extracts were prepared by using the same lysis buffer as for DNA pull-down assay. Cell lysates were homogenized and cleared. Cleared lysates were incubated with IgG or anti-V5 overnight at 4°C. Protein A Sepharose were added to lysates and incubated for 1-2 hr at 4°C. Bound proteins were washed extensively with lysis buffer. Bound DNAs were eluted by RNase and Proteinase K treatment. Eluted DNAs were extracted by phenol-chloroform extraction and purified by ethanol precipitation. Purified DNAs were run on a 6% Urea-denaturing gel, stained with SYBR gold (Invitrogen), and analyzed.

### Colony formation assay

To examine colony forming ability of ABCF1 overexpressing cells under DNA damaging condition, D3 mouse ES cells were transduced with lentiviruses expressing RFP or full-length human ABCF1 (pHAGE-RFP and pHAGE-V5-hABCF1). Mouse ES cells were transduced with lentiviruses at MOI 3 and selected with neomycin (500 *μ*g/ml). 200 RFP or ABCF1 overexpressing mouse ES cells were plated on 24-well plates. Cells were allowed to recover for 24 hr before treatment with DMSO or ETO (1 *μ*M) for 1, 2, or 4 hr, after which fresh medium without ETO was replaced. After 6 days, cells were fixed and stained for AP activity (EMD Millipore). AP-positive cells were counted and analyzed.

### Amino acid composition analysis

The amino acid composition of a protein was analyzed in R with custom scripts. The occurrence of each amino acid is counted by using the package ‘stringr’ and plotted with the package ‘plot. matrix.

### Statistical analysis

To determine statistical significance, *P*-values were calculated by using unpaired two-sided Student’s t-test. *P*-values less than 0.05 (<0.05) were considered as statistically significant and they were indicated with * (\**P*<0.05). All data represent the mean ± SEM (error bars) except for figure 7F. For figure 7F, box-and-whisker plot was used. The central mark represents median and edges represent 25^th^ and 75^th^ percentiles. The whiskers indicate 5^th^ to 95^th^ percentiles.

## ACKNOWLEDGEMENTS

We thank G. Dailey for help with expression constructs; S. Zheng for tissue culture assistance; and S. Zhou and D. King for mass spectrometry analysis. We thank R. Tjian, Y-C. Hsu, S. Agarwal, Y. Isogai, Z. Zhang, B. Zhang, E. Guo, S. Chong, and C. Cattoglio for valuable discussion. We acknowledge the following funding sources: NIH grant R01HL125527, Harvard Stem Cell Institute, Charles H. Hood Foundation, and Brigham and Women’s Hospital HVC Junior Faculty Research Awards to Y.W.F.

## SUPPLEMENTARY FIGURES AND TABLES

**Supplementary Figure S1.**
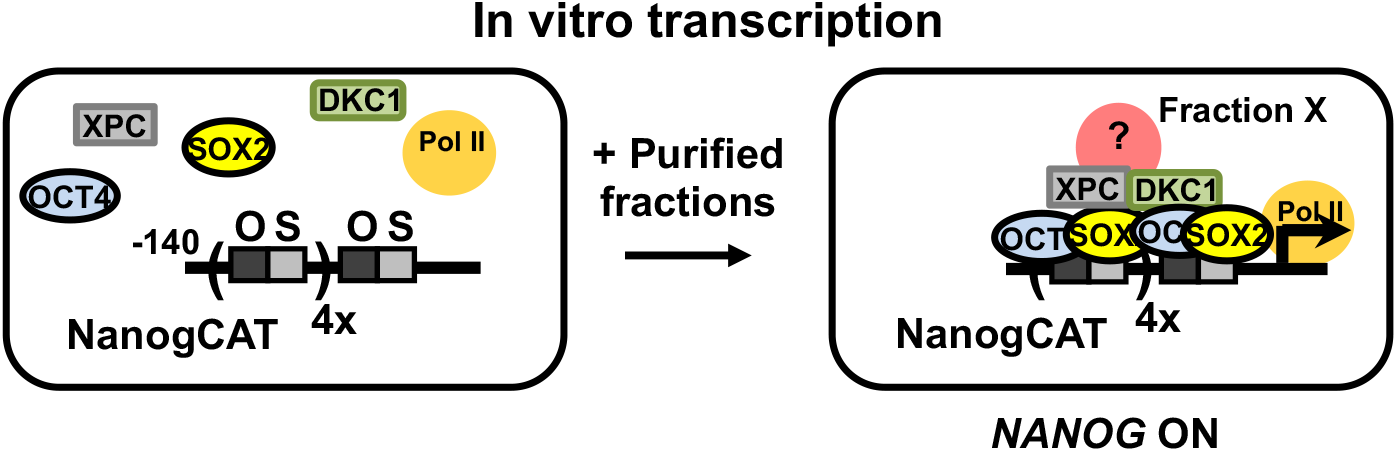
Schematic diagram of in vitro transcription assay to detect SCC-B activity. In vitro transcription reactions contain purified general transcription factors (TFII-A, B, D, E, F, and H), RNA polymerase II (Pol II), recombinant XPC and DKC1 complexes, and affinity-purified OCT4 and SOX2. These factors are supplemented with partially purified fractions from NT2 nuclear extracts and assayed for the ability of protein fractions to stimulate transcription of a modified transcription template derived from the human *NANOG* promoter (−140). The modified *NANOG* template contains four extra copies of the OCT4/SOX2 composite binding sites upstream of the native enhancer. We have previously shown that SCC-dependence transcriptional activation is also observed using an unadulterated, native *NANOG* promoter template (Fong et al., 2011). Transcription is measured by primer extension assay.

**Supplementary Figure S2.**
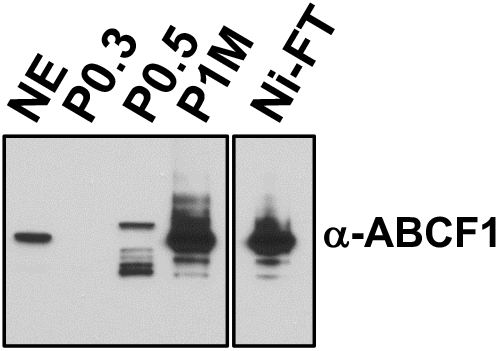
ABCF1 is highly enriched in the transcriptionally active, partially purified nuclear extract fractions. Comparative western blot analysis of NT2 nuclear extract (NE), phosphocellulose 0.3 M, 0.5 M, 1 M KCl fractions (P0.3, P0.5, P1M, respectively), and Ni-NTA flowthrough (Ni-FT) using an antibody against ABCF1. P0.3 and P0.5 fractions do not contain SCCs (Fong et al., 2011). Each lane contains 5 *μ*g of protein as determined by Bradford protein assay.

**Supplementary Figure S3.**
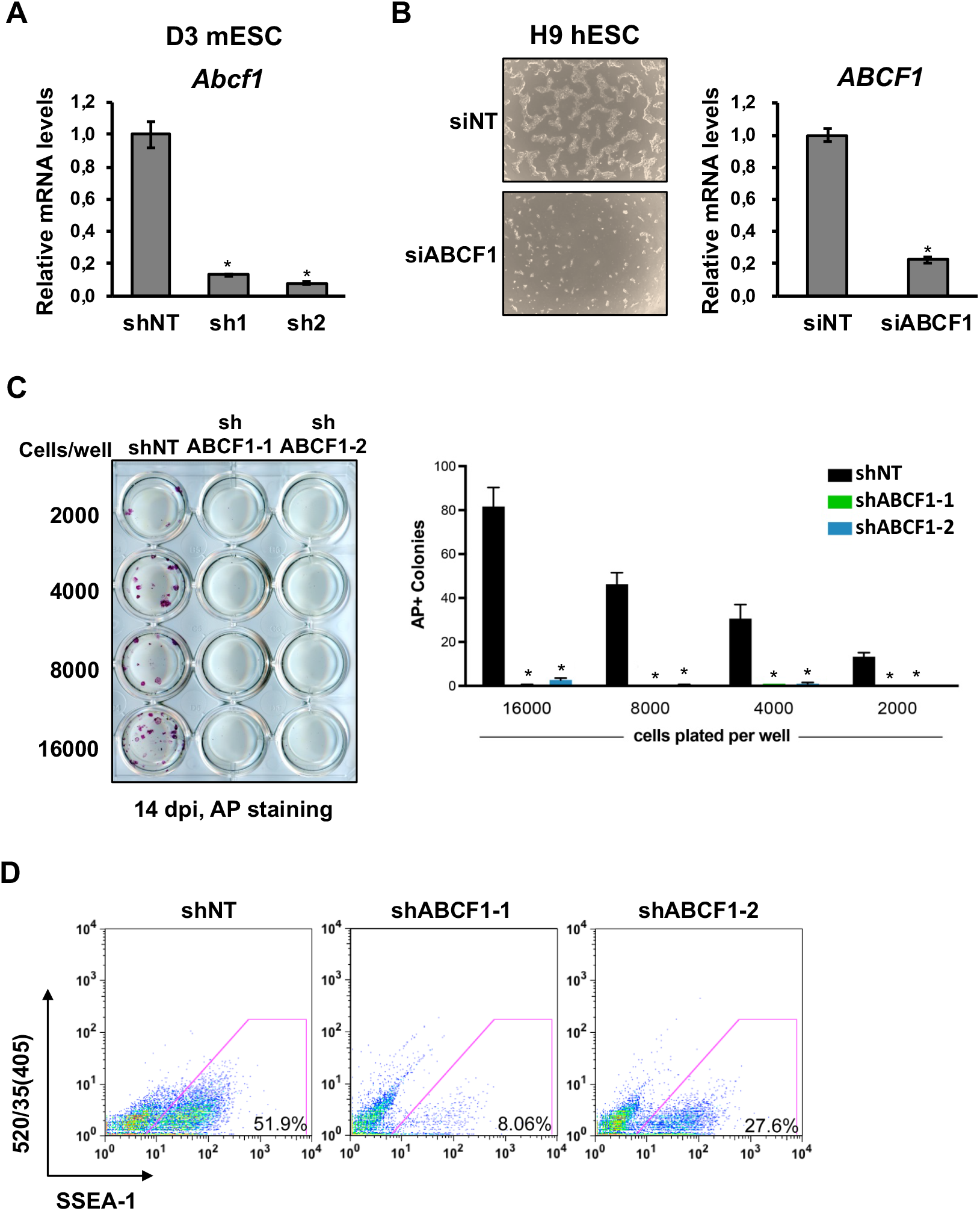
(A) Efficiency of shRNA-mediated knockdown of ABCF1 in D3 mouse ES cells is determined by qPCR. *Abcf1* mRNA levels are normalized to *Actb* and expressed as fraction of control (=1). (B) H9 human ES cells are transfected with non-targeting siRNA (siNT) or siRNAs against ABCF1 (siABCF1). Knockdown efficiency is analyzed by qPCR as in (A). Downregulation of ABCF1 in human ES cells compromises stem cell self-renewal. Phase contrast images of human H9 ES cells transfected with siNT or siABCF1. (C) Depletion of ABCF1 blocks somatic cell reprogramming. CF-1 MEFs are transduced with lentiviruses expressing either a control shRNA (NT) or shRNAs targeting ABCF1 (shABCF1-1 and shABCF1-2), together with lentiviruses expressing OCT4, KLF4, SOX2, and c-MYC. Cells are plated at the indicated number in 24-well plates; cellular reprogramming is initiated by the addition of doxycycline (dox). Cells are stained for alkaline phosphatase (AP) activity (left) and counted (right) after 14 days (11 days with dox followed by 3 days of dox withdrawal) post induction (dpi). (D) Single cells suspensions of 14 dpi reprogrammed CF-1 MEFs as described in (C) are stained with anti-mouse SSEA-1 and analyzed by flow cytometry. Error bars present SEM. *n* = 3. (*) *P* < 0.05, calculated by two-sided Student’s t-test.

**Supplementary Figure S4.**
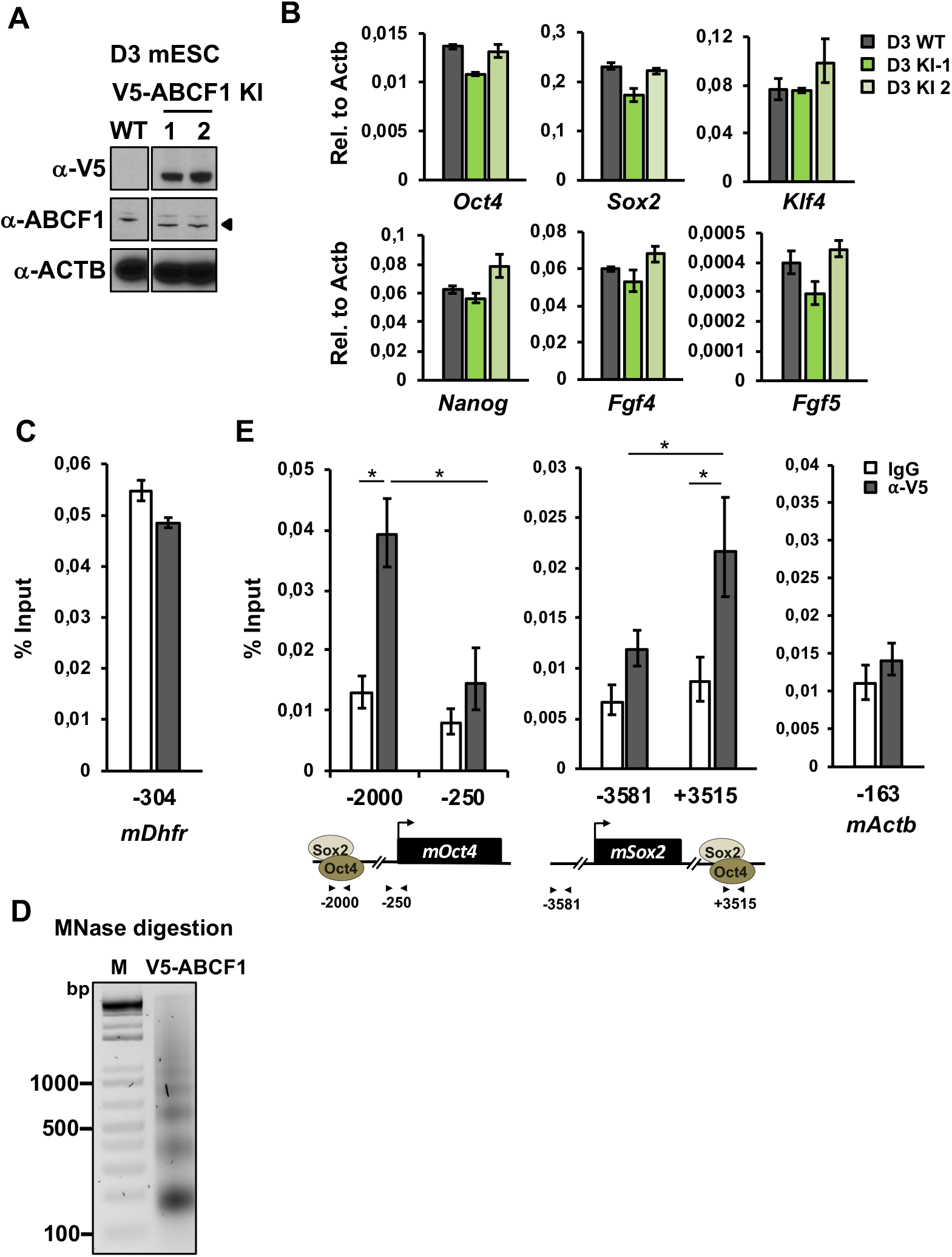
(A) Expression of V5-tagged ABCF1 (V5-ABCF1) in two independent knock-in (KI) D3 mouse ES cell clones are verified by western blotting using antibodies against V5 and ABCF1. ACTB is used as loading control. (B) V5 tagging of ABCF1 in mouse ES cells (D3 KI-1, −2) does not compromise ABCF1 function. mRNA levels of *Oct4*, *Sox2*, *Nanog*, *Klf4*, and *Fgf4* are quantified by qPCR and normalized to *Actb*. V5-ABCF1 knock-in clones display low expression level of differentiation-associated gene, *Fgf5*, comparable to WT cells. (C) ChIP-qPCR analysis of ABCF1 enrichment on housekeeping gene *Dhfr* as described in Figure 4. (D) MNase-digested chromatin from EGS/formaldehyde crosslinked V5-ABCF1 KI mouse ES cells are purified and separated on an agarose gel. (E) MNase-ChIP analysis of ABCF1 on the *Oct4, Sox2,* and *Actb* genes by qPCR as described in Figure 4. Error bars present SEM. *n* = 3. (*) *P* < 0.05, calculated by two-sided Student’s t-test.

**Supplementary Figure S5.**
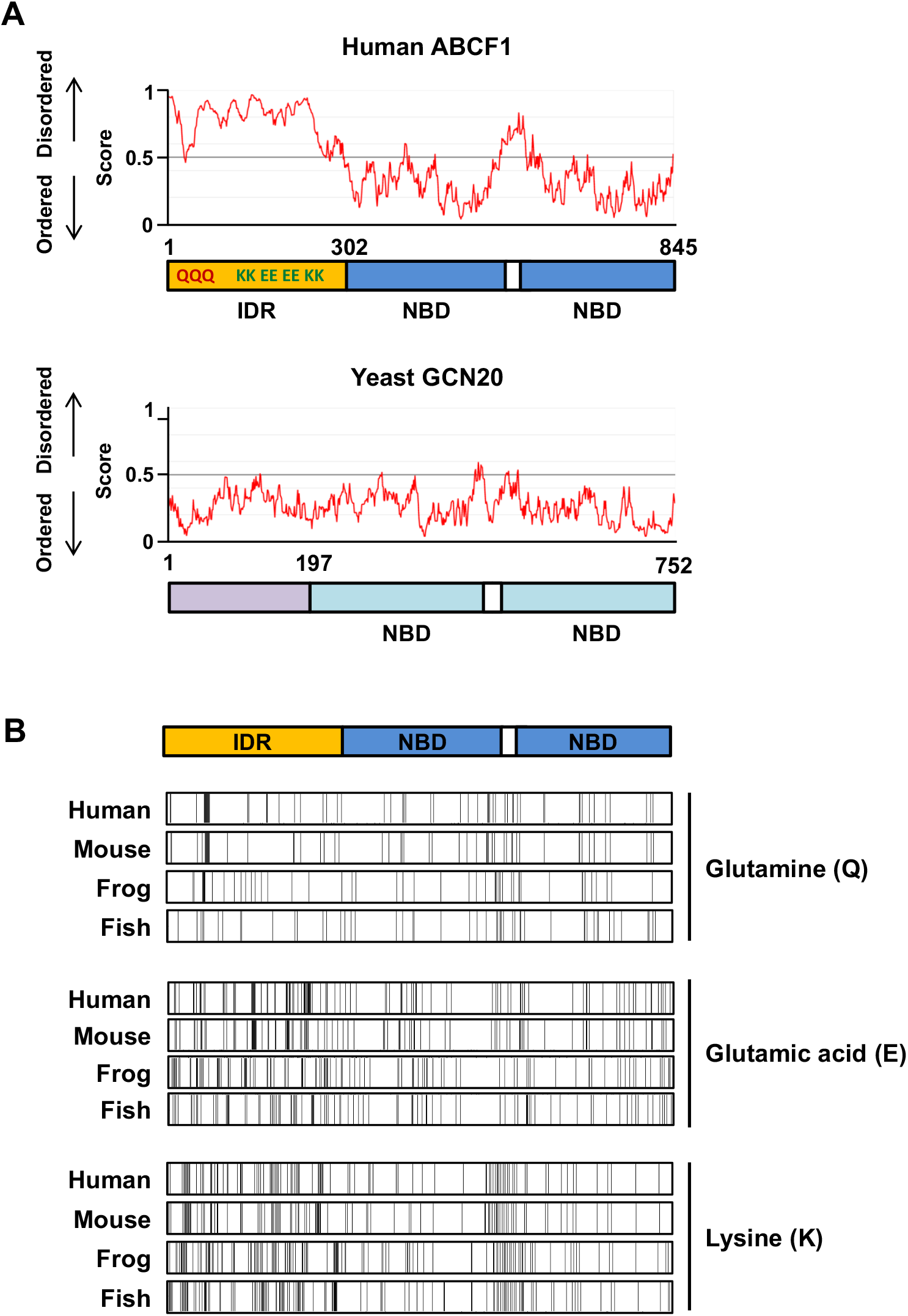
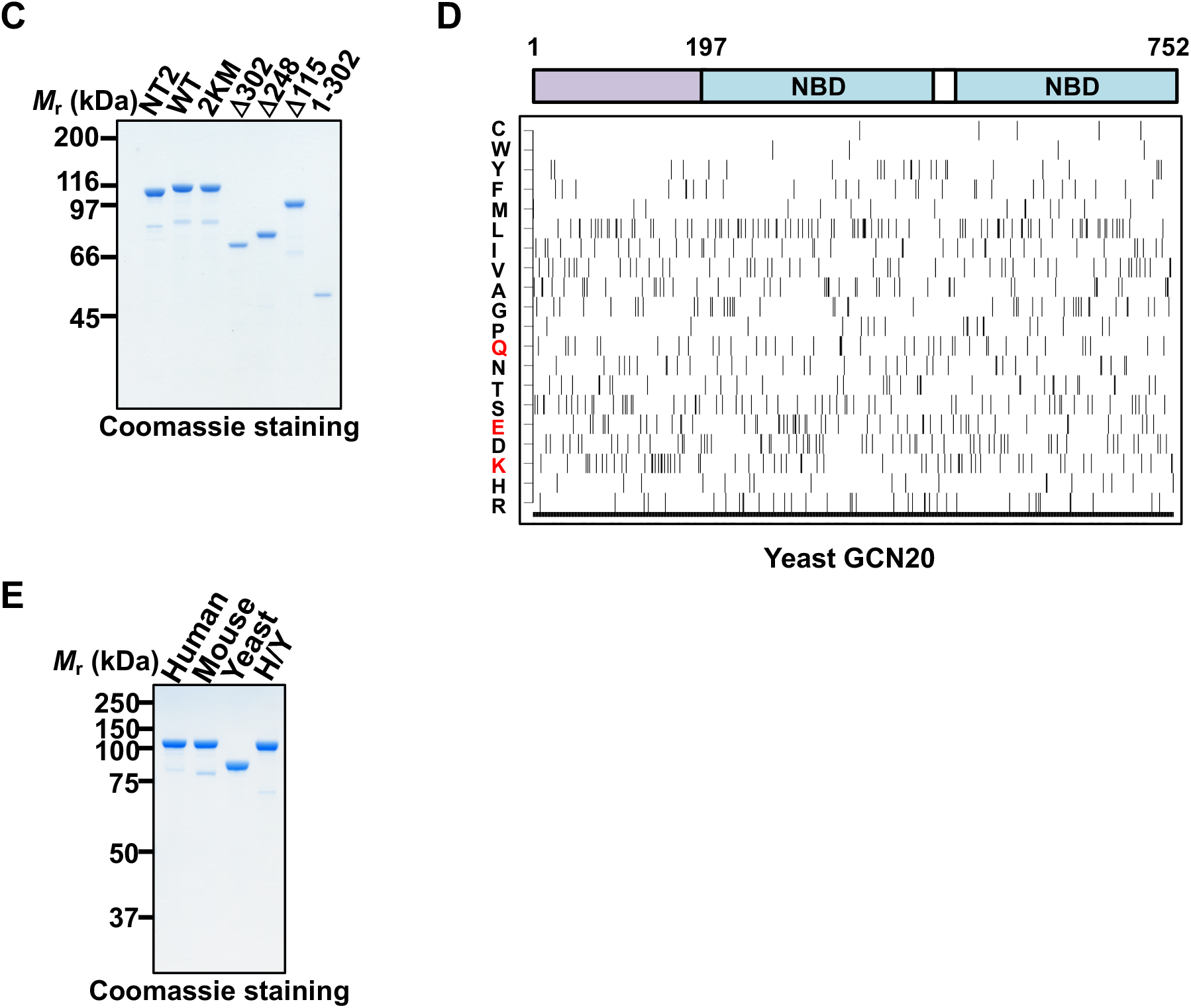
(A) Intrinsically disordered protein prediction tool, IUPred2 (https://iupred2a.elte.hu/) (Abor et al., 2018), is used to identify unstructured regions in both human ABCF1 and yeast GCN20. X-axis indicates position of the amino acids and Y-axis shows the probability of disordered sequences. Regions that are above the value of 0.5 are predicted to be unstructured and those below 0.5 are more likely structured. Relative positions of the protein domains are depicted below. Similar results are also obtained using a different prediction program DISOPRED3 (data not shown) (Jones and Cozzetto, 2015). (B) Comparison of distribution of glutamine (Q), glutamic acid (E), and lysine (K) residues in ABCF1 from indicated organisms, analyzed as described in Figure 5A. (C) SDS-PAGE and Coomassie staining of endogenous ABCF1 purified from NT2 nuclear extracts (NT2), purified recombinant full-length and truncated ABCF1 proteins described in Figure 5B. (D) Amino acid composition of yeast GCN20, analyzed as described in Figure 5A. The relative positions of NBDs and the non-conserved N-terminal region of GCN20 are indicated. One-letter abbreviations for amino acids are indicated (Left): C, Cys; W, Trp; Y, Tyr; F, Phe; M, Met; L, Leu; I, Ile; V, Val; A, Ala; G, Gly; P, Pro; Q, Gln; N, Asn; T, Thr; S, Ser; E, Glu; D, Asp; K, Lys; H, His; R, Arg. (E) SDS-PAGE and Coomassie staining of recombinant human, mouse, and yeast homolog of ABCF1 (GCN20) as well as the human-yeast hybrid protein (H/Y).

**Supplementary Figure S6.**
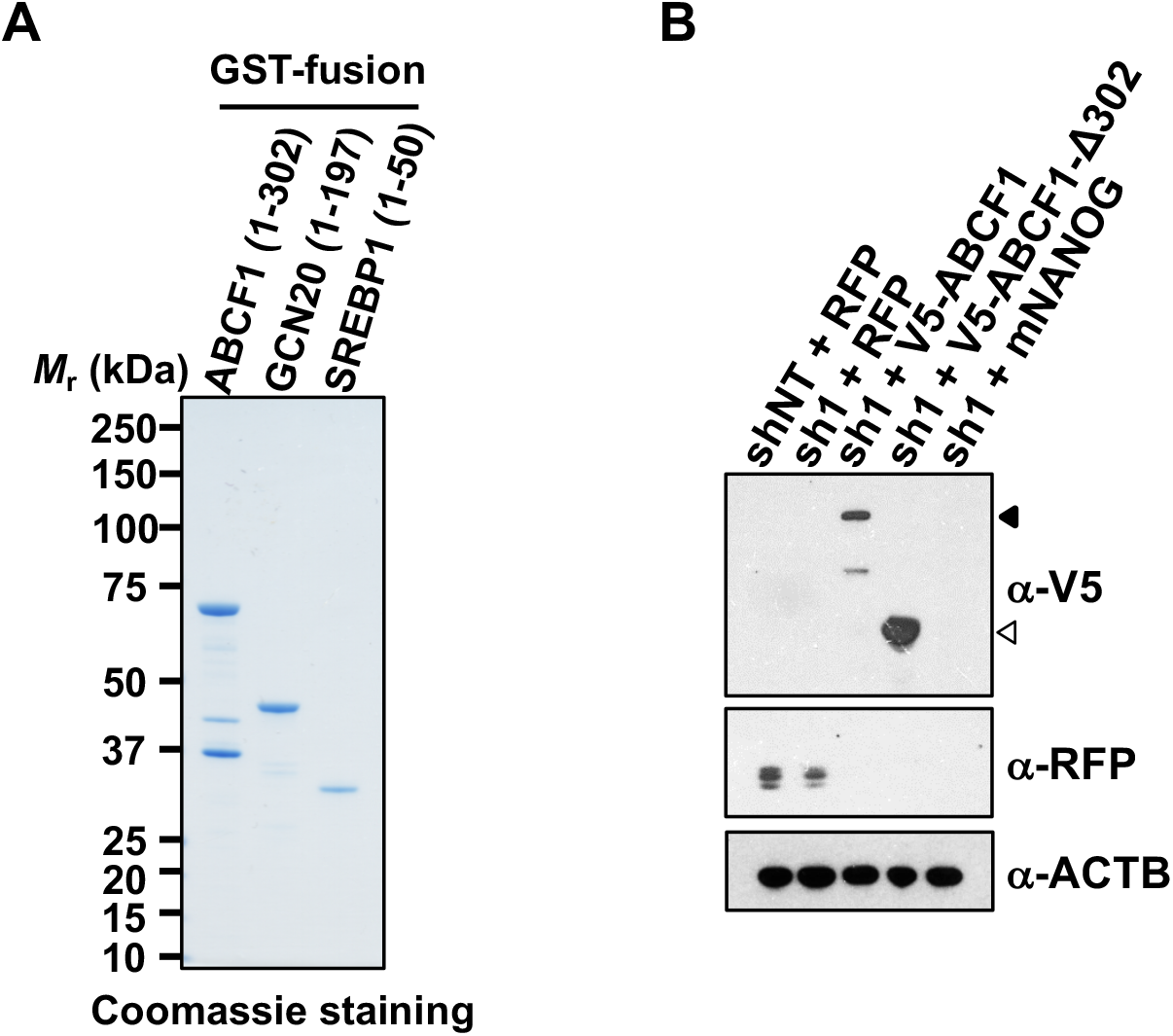
(A) SDS-PAGE and Coomassie staining of purified GST-fusion proteins containing N-terminal domain of ABCF1 (1-302), GCN20 (1-197), and SREBP1a (1-50). Comparison of distribution of glutamine (Q), glutamic acid (E), and lysine (K) residues in ABCF1 from indicated organisms, analyzed as described in Figure 5A. (B) Expression of control RFP and V5-tagged human ABCF1 proteins described in Figure 6E is confirmed by western blotting using antibodies against RFP and V5. Filled and white arrowheads denote V5-tagged full-length and IDR-truncated human ABCF1, respectively. ACTB is used as loading control.

**Supplementary Figure S7.**
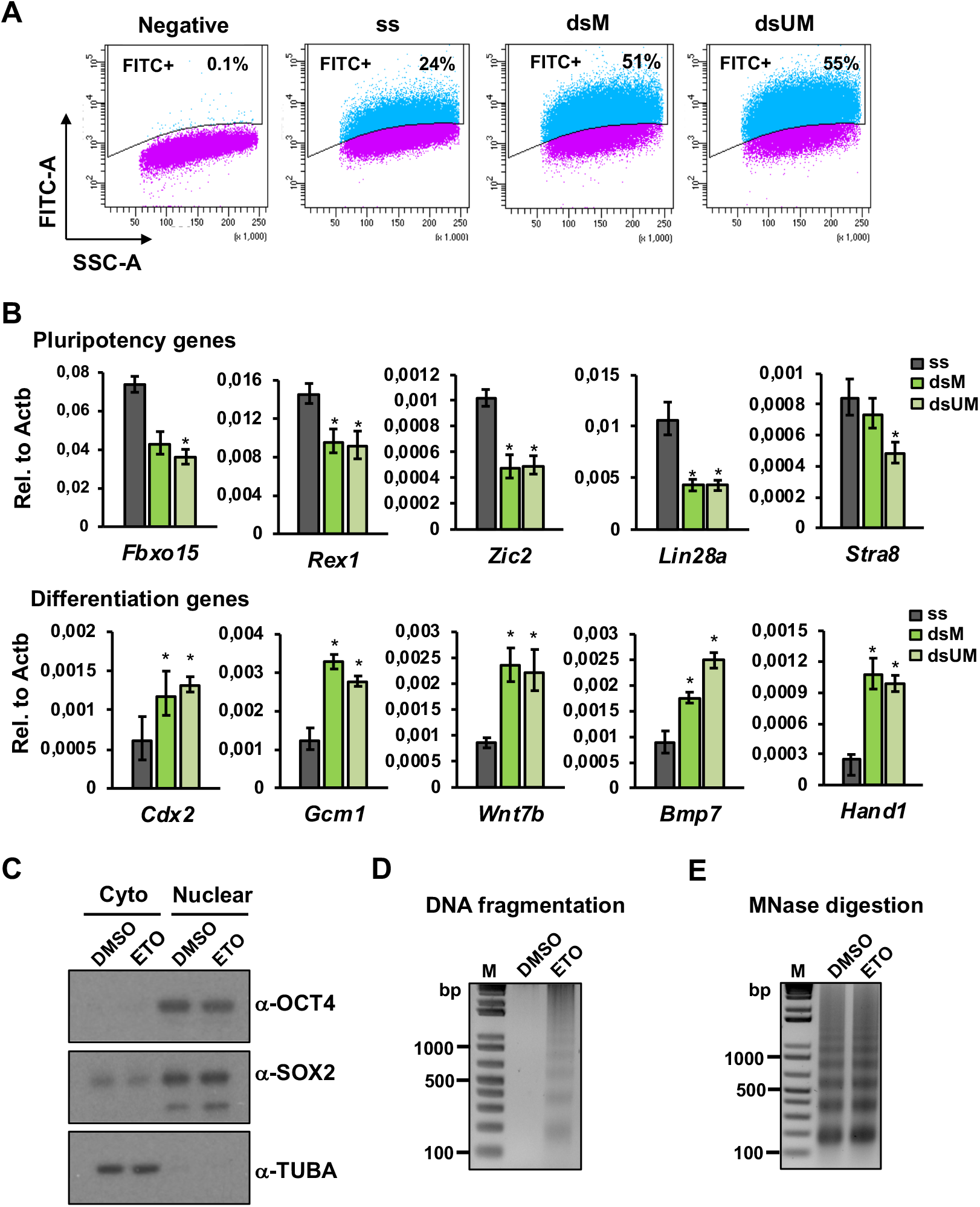
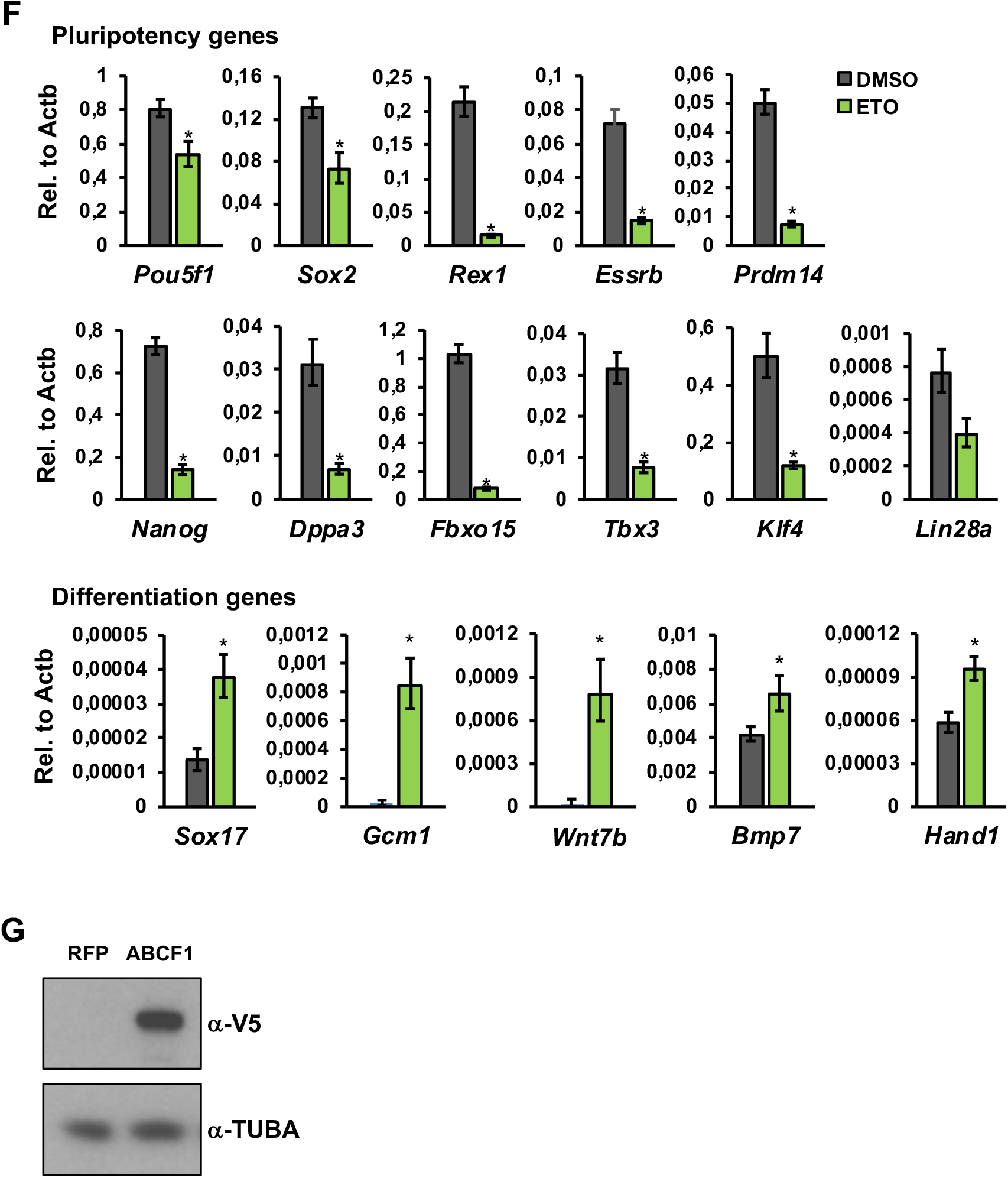
(A) Single cell suspensions of D3 mouse ES cells transfected with 5’ 6-carboxyfluorescein (6-FAM) labeled ss, ds-M, or ds-UM on the sense strand are analyzed by flow cytometry. 6-FAM-positive cells (blue) are gated by comparing to untransfected cells (Negative). 6-FAM-positive cells are purified for further analyses. (B) Expression levels of pluripotency and differentiation genes in mouse ES cells as sorted in (A) are analyzed by qPCR. (C) Western blot analysis of nuclear and cytoplasmic fractions prepared from V5-ABCF1 knock-in mouse ES cells using antibodies against OCT4, SOX2, and TUBA. Effective nuclear-cytoplasmic fractionation is demonstrated by enrichment of OCT4 and SOX2 in nuclear extracts and TUBA in cytoplasmic fraction. (D) DNAs purified from DMSO or ETO-treated (80 μM) V5-ABCF1 KI mouse ES cells grown in 2i/LIF medium are separated on an agarose gel and visualized by ethidium bromide staining. (E) Crosslinked nuclear chromatin from DMSO or ETO-treated (80 μM) mouse cells are fragmented by MNase digestion. Digested DNAs are purified, separated on an agarose gel, and stained with ethidium bromide. Both DMSO and ETO-treated chromatins are digested to a similar degree. (F) Expression levels of pluripotency and differentiation genes in DMSO or ETO-treated (80 μM) V5-ABCF1 KI mouse ES cells are analyzed by qPCR, normalized to *Actb*. Error bars present SEM. *n* = 3. (*) *P* < 0.05, calculated by two-sided Student’s t-test. (G) Western blot analysis showing stable overexpression of V5-ABCF1 in wild-type D3 mouse ES cells transduced with lentiviruses expressing V5-ABCF1 compared to control mouse ES cells expressing RFP.

**Supplementary Table S1.**
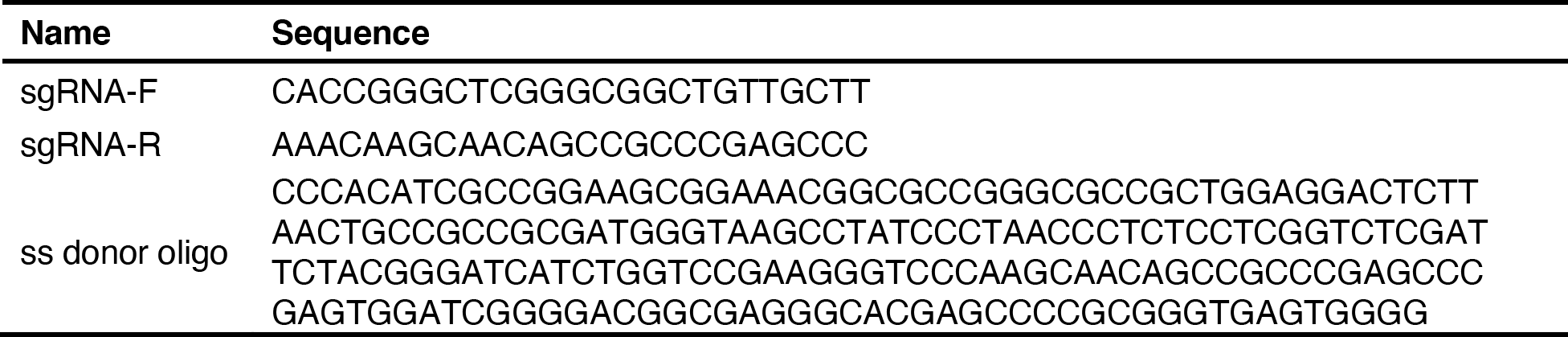
sgRNA and ss donor oligo sequences for generation endogenously V5-tagged ABCF1 knock-in mouse ES cell line.

**Supplementary Table S2.**
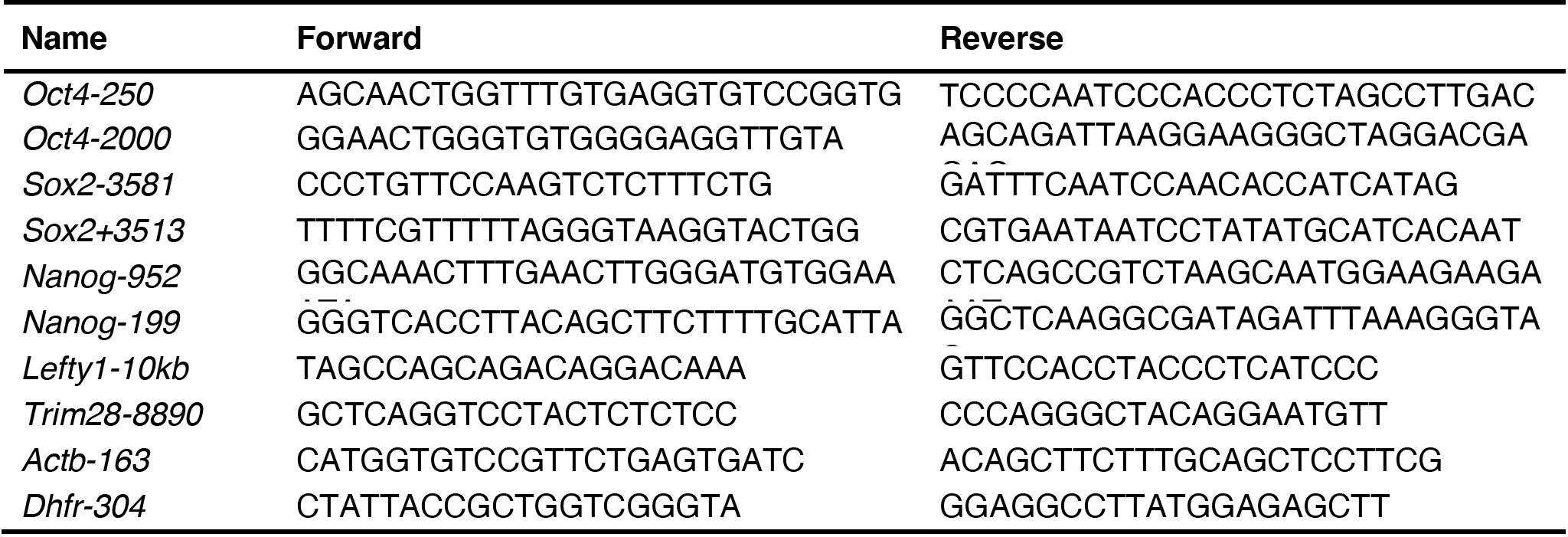
Primers used for ChIP or MNase ChIP-qPCR analysis.

**Supplementary Table S3.**
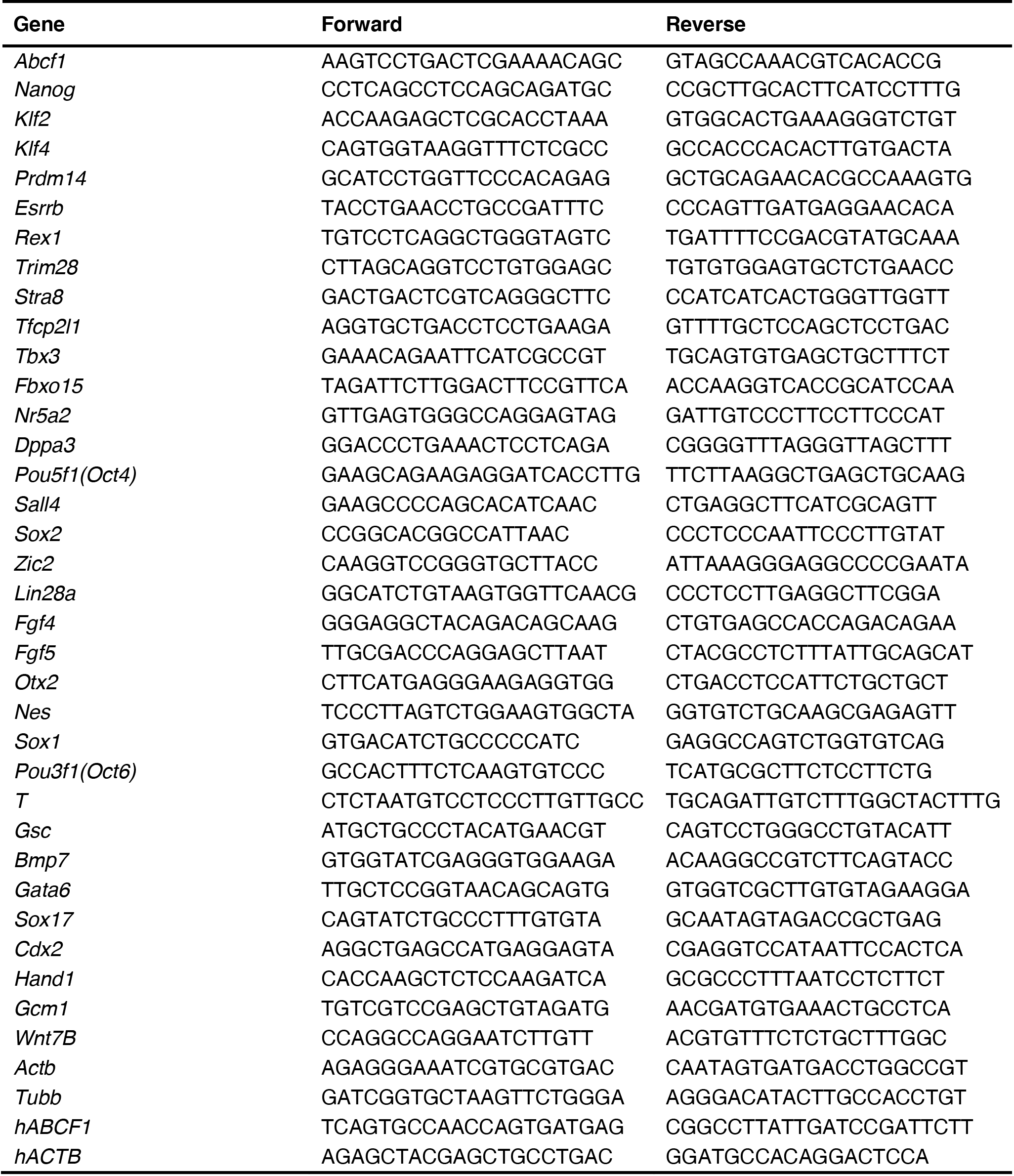
Primers used for RT-qPCR analysis.

**Supplementary Table S4.**
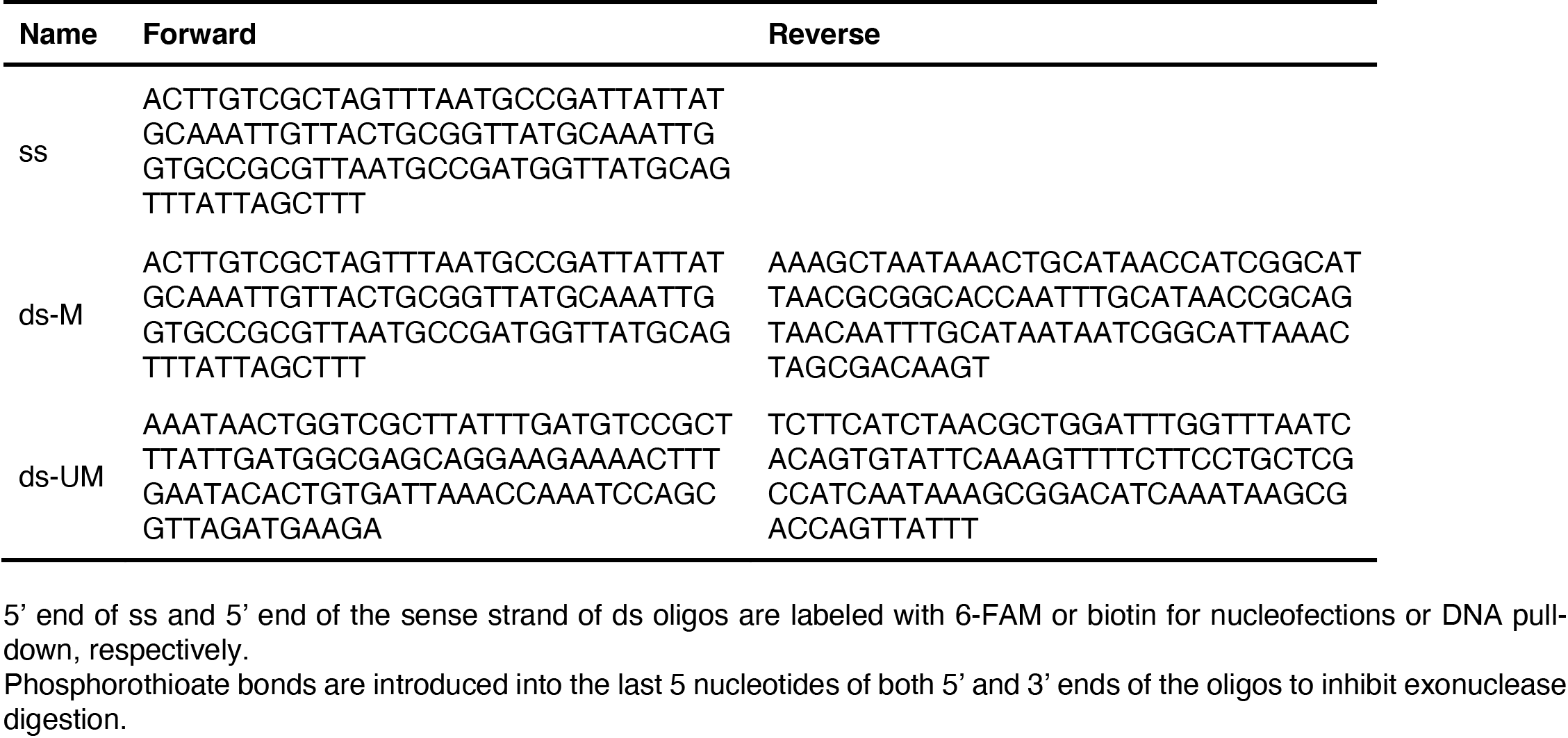
DNA oligos used for nucleofections and DNA pull-down assays.

## Notes

### Competing Interest Statement

The authors have declared no competing interest.

### Summary of Updates

Due to the following message from the journal we submitted: Per the journal policy, only the original submission may be posted on not-for-profit preprint servers. Please post your original submission (dated November 15) over the current posting, so that the landing page goes directly to that original rather than to the revised version. Note that we cannot move your manuscript forward to the handling editor until this issue resolved.

